# Profiling the genome-wide landscape of tandem repeat expansions

**DOI:** 10.1101/361162

**Authors:** Nima Mousavi, Sharona Shleizer-Burko, Richard Yanicky, Melissa Gymrek

## Abstract

Tandem Repeat (TR) expansions have been implicated in dozens of genetic diseases, including Huntington’s Disease, Fragile X Syndrome, and hereditary ataxias. Furthermore, TRs have recently been implicated in a range of complex traits, including gene expression and cancer risk. While the human genome harbors hundreds of thousands of TRs, analysis of TR expansions has been mainly limited to known pathogenic loci. A major challenge is that expanded repeats are beyond the read length of most next-generation sequencing (NGS) datasets and are not profiled by existing genome-wide tools. We present GangSTR, a novel algorithm for genome-wide genotyping of both short and expanded TRs. GangSTR extracts information from paired-end reads into a unified model to estimate maximum likelihood TR lengths. We validate GangSTR on real and simulated data and show that GangSTR outperforms alternative methods in both accuracy and speed. We apply GangSTR to a deeply sequenced trio to profile the landscape of TR expansions in a healthy family and validate novel expansions using orthogonal technologies. Our analysis reveals that healthy individuals harbor dozens of long TR alleles not captured by current genome-wide methods. GangSTR will likely enable discovery of novel disease-associated variants not currently accessible from NGS.

## 1 Introduction

Next-generation sequencing (NGS) has the potential to profile nearly all genetic variants simultaneously in a single assay. Indeed, whole exome sequencing (WES) and whole genome sequencing (WGS) have successfully identified single nucleotide polymorphisms (SNPs) and small indels contributing to a range of phenotypes, including Mendelian diseases [47], cancer [3], and complex traits [5]. Recently, several studies have demonstrated the power of NGS to genotype more complex structural variants (SVs) and revealed a contribution to a variety of traits including gene expression [8], cancer [45], and autism spectrum disorder [7]. Despite this progress, NGS pipelines struggle with highly repetitive regions of the genome, which are still routinely filtered from most studies.

Here, we focus on short tandem repeats (STRs) with motif lengths of 1-6bp and variable number tandem repeats (VNTRs) with motif lengths of up to 20bp, which we collectively refer to as TRs. TRs have been implicated in dozens of disorders [29], such as Huntington’s Disease and Fragile X Syndrome, which together affect millions of individuals worldwide [17, 33, 35]. In most cases, the pathogenic mutation is an expansion of the number of repeats. Importantly, known pathogenic TRs represent just a small fraction of the more than one million TRs in the human genome [43]. Recently, thousands of TRs have been shown to play a role in gene regulation [13, 34] and it is becoming increasingly clear that TRs across the genome are likely to have widespread contributions to complex polygenic traits [32, 14, 15]. In these cases, smaller expansions or contractions may subtly increase or decrease risk for a trait, similar to effect sizes observed for point mutations, and work together to modulate an individual’s disease risk [41].

Over the last several years, we and others have developed a series of tools for genome-wide genotyping of STRs [12, 44, 16, 23] from short reads or targeted genotyping of VNTRs [4] from both short and long reads. These tools primarily rely on identifying reads that completely enclose the repeat of interest. While most TRs in the human genome can theoretically be spanned by 100bp reads [27], in practice repeats longer than around 70bp are difficult or impossible to genotype due to an insufficient number of enclosing reads. Notably, in our recent genome-wide analysis [36] using HipSTR [44], more than 150,000 STRs were filtered because they showed strong departure from genotype frequencies expected Hardy-Weinberg equilibrium, in part because of dropout of long alleles. The list of filtered TRs includes most known pathogenic TR expansions, for which even normal alleles typically exceed the length of short reads [15]. Thus existing NGS pipelines provide an incomplete picture of genome-wide variation at TRs.

Recently, several methods have been developed to analyze expanded TRs from NGS, but all face limitations that do not allow for unbiased genome-wide analysis of TR lengths. exSTRa [39] classifies a repeat as “expanded” vs. “normal” but requires a control cohort and does not estimate repeat length, which is often informative of disease severity or age of onset [15] and is required for performing genome-wide association studies. STRetch [9] can perform genome-wide expansion identification but does not analyze short TRs, is limited to motifs of up to 6bp, and is computationally expensive. Tredparse [38] models multiple aspects of paired end reads but cannot estimate repeat lengths longer than the sequencing fragment length. ExpansionHunter [11] produces accurate genotypes across a range of repeat lengths except when both alleles are close to or longer than the sequencing read length. Finally, Tredparse and ExpansionHunter have been primarily designed for targeted analysis of known pathogenic expansions and do not scale genome-wide.

Long read technologies have recently been applied to genotype long and complex repeats, such as the CCG repeat implicated in Fragile X Syndrome [26] and a complex pentamer repeat implicated in myoclonus epilepsy [18]. While long reads offer a potential solution to genome-wide TR analysis, NGS remains the gold standard for diagnostic sequencing and population-wide studies due to its low cost and substantially higher throughput [30]. Furthermore, the low per-base accuracy and high indel rate of long read technologies present major challenges to accurate quantification of repeat counts, especially for TRs with short motif lengths. Thus, we focus here on the challenge of comprehensive TR genotyping from short reads.

Here, we present GangSTR, a novel method for genome-wide analysis of TRs from NGS data. GangSTR relies on a general statistical model incorporating multiple properties of paired-end reads into a single maximum likelihood framework capable of genotyping both normal length and expanded repeats. We extensively benchmark GangSTR against existing methods on both simulated and real datasets harboring a range of allele lengths and show that GangSTR is both faster and more accurate than existing solutions. Finally, we apply GangSTR to genotype TRs using high-coverage NGS from a trio family to evaluate Mendelian inheritance and validate novel repeat expansions using orthogonal long read and capillary electrophoresis data. Altogether, our analyses demonstrate GangSTR’s ability to robustly genotype a range of TR classes, which will likely enable identification of novel pathogenic expansions as well as genome-wide association studies of TR variation in large cohorts.

GangSTR is packaged as an open-source tool at https://github.com/gymreklab/GangSTR.

## 2 MATERIALS AND METHODS

### 2.1 Overview of the GangSTR model

GangSTR is an end-to-end method that takes sequence alignments and a reference set of TRs as input and outputs estimated diploid repeat lengths. Its core component is a maximum likelihood framework incorporating various sources of information from short paired-end reads into a single model that is applied separately to each TR in the genome.

Multiple aspects of paired-end short reads can be informative of the length of a repetitive region. Reads that completely enclose a repeat trivially allow determination of the repeat number by simply counting the observed number of repeats. While most of existing tools have primarily focused on repeat-enclosing reads, other pieces of information, such as fragment length, coverage, and existence of partially enclosing reads, are all functions of repeat number. Recent tools for targeted genotyping of expanded STRs utilize various combinations of these information sources (Table 1).

**Table 1:** Classes of read pairs and features used by existing tools for genotyping TRs from short reads.

| Tool | Enclosing | FRR | Spanning | Off-target FRR | Estimates # rpt. | Genome-wide | Estimation limit |
| --- | --- | --- | --- | --- | --- | --- | --- |
| LobSTR [12] | X |  |  |  | X | X | < Read Length |
| HipSTR [44] | X |  |  |  | X | X | < Read Length |
| STRetch [9] |  | X |  | X | X | X | Only reports expanded TRs |
| exSTRa [40] |  | X |  | X |  | X | Does not estimate TR length |
| Tredparse [38] | X | X | X |  | X |  | < Fragment Length |
| ExpansionHunter [11] | X | X |  | X |  | X | Poor performance when both alleles long |
| GangSTR | X | X | X | X | X | X | Not limited by fragment or read length |

GangSTR incorporates each of these informative aspects of paired-end read alignments into a single joint likelihood framework (Figure 1). The underlying genotype is represented as a tuple 〈*A, B*〈, where *A* and *B* are the repeat lengths of the two alleles of an individual. We define four classes of paired-end reads: enclosing read pairs (“E”) consist of at least one read that contains the entire TR plus non-repetitive flanking region on either end; spanning read pairs (“S”) originate from a fragment that completely spans the TR, such that each read in the pair maps on either end of the repeat; flanking read pairs (“F”) contain a read that partially extends into the repetitive sequence of a read; and fully repetitive read pairs (“FRR”) contain at least one read consisting entirely of the TR motif. Two types of probabilities are computed for each read pair: the class probability, which is the probability of observing a read pair of a given class given the true genotype, and the read probability, which gives the probability of observing a particular characteristic of the read pair. A different characteristic is modeled for each class (Figure 2).

**Figure 1:**
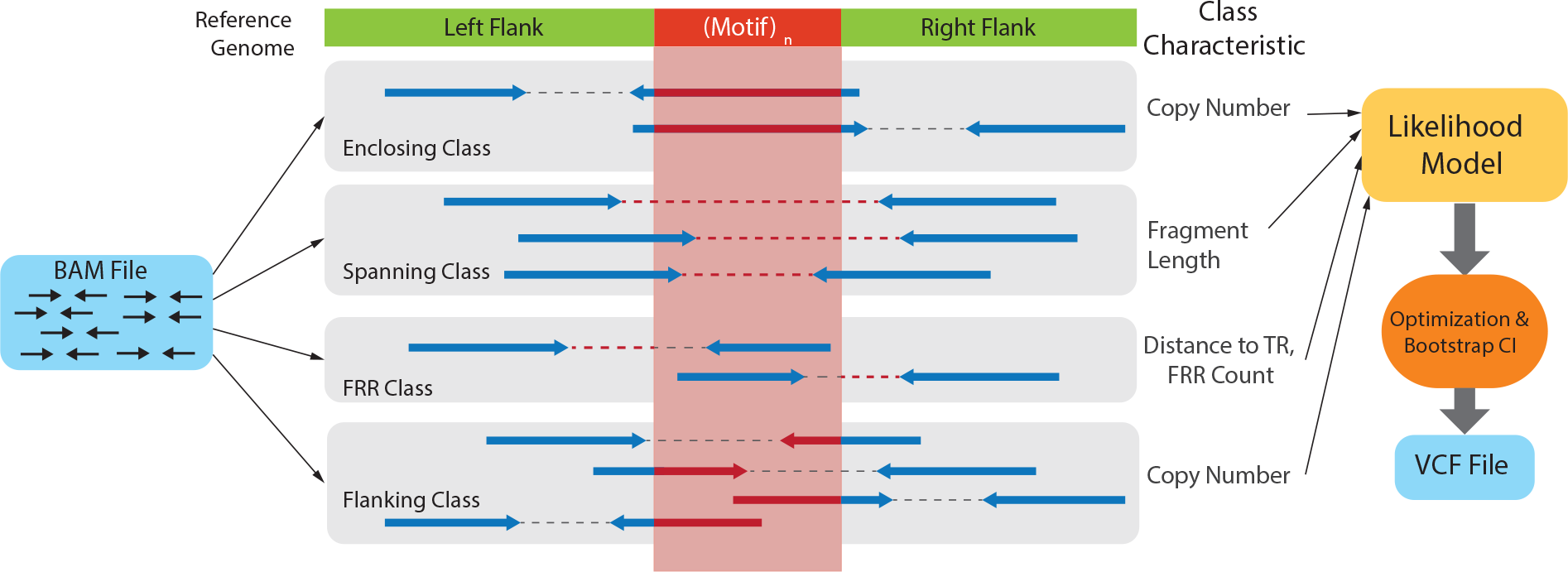
Schematic of GangSTR method. Paired end reads from an input set of alignments are separated into various read classes, each of which provides information about the length of the TR in the region. This information is used to find the maximum likelihood diploid genotype and confidence interval on the repeat length. Results are reported in a VCF file.

**Figure 2:**
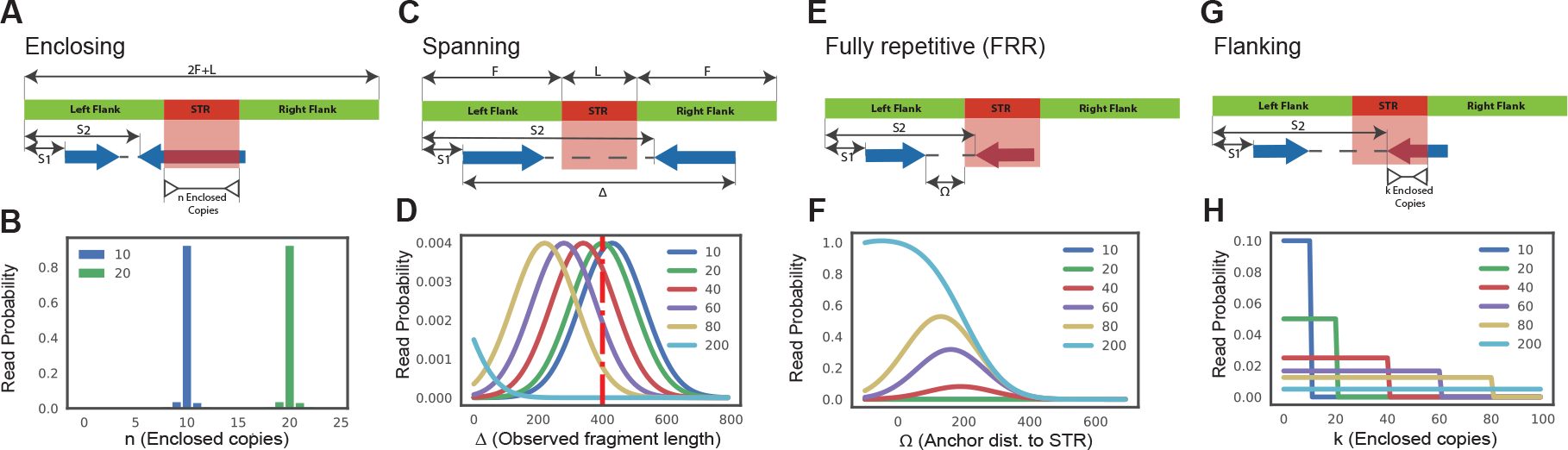
Four classes of informative read pairs. A. Enclosing class: characteristic *n* corresponds to the number of repeat copies enclosed in the read. B. *n* is modeled for different repeat length accounting for errors introduced during PCR. C. Spanning class: characteristic ∆ denotes the observed fragment length for a read pair. D. ∆ is modeled for different repeat lengths. Longer repeats give shorter observed fragment lengths. The red vertical dashed line gives the mean actual fragment length. E. Fully Repetitive Read (FRR) class: characteristic Ω is the distance of the non-repetitive read from the repeat region. F. Ω is modeled for different repeat lengths. Longer repeats give shorter observed Ω values. G. Flanking class: characteristic *k* shows the number of copies extracted from the flanking read. H. *k* is modeled for different repeat lengths. *S*1 and *S*2 give the start coordinates of each read in the pair relative to the beginning of the first flanking region. For A, C, E, and G, *F* shows the length (bp) of the flanking region and the repeat is *L* bp long (*A* copies of a repeat of length *m*). For B, D, F, and H, each color denotes a different repeat length (blue=10 copies, green=20 copies, red=40 copies, purple=60 copies, gold=80 copies, light blue=200 copies).

### 2.2 Computation of Log Likelihood

The likelihood model computes the probability of the observed read pairs given a true underlying diploid genotype:

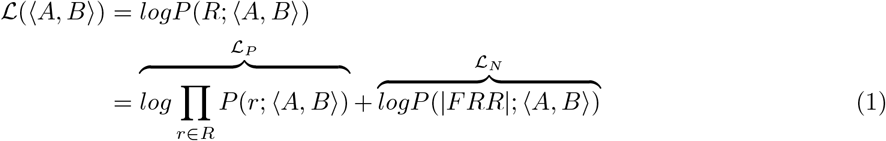

Where 𝓛(〈*A, B*〉) corresponds to the total log likelihood of genotype 〈*A, B*〉, which consists of term 𝓛_*P*_ combining the contribution of each read pair *r* from the set of informative read pairs *R*, and term 𝓛_*N*_ which models the total number of FRR reads.

#### 2.2.1. Read pair term

The first term in (1) is calculated by extracting characteristics from every informative read pair, where the specific characteristic modeled depends on the class of the read. Each read pair is assigned to one or more classes. If a read pair belongs to multiple classes (for example, a read pair can be both spanning and flanking), it appears once in each class for its contribution to both likelihood classes. The read pair term is computed as follows:

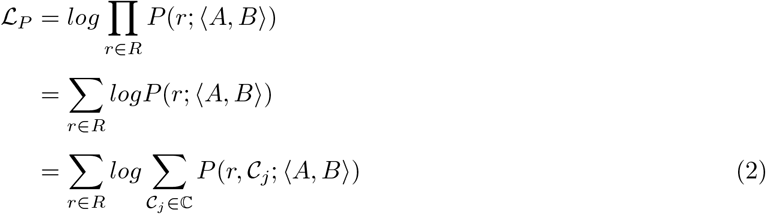

where 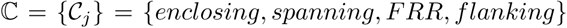 is the set of all informative read classes. Every informative read pair *r* belongs to a class of informative reads, we denote this class by *C*(*r*). The value of 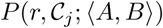 is set to 1 if 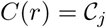 and 0 otherwise. We thus simplify the term for each read pair:

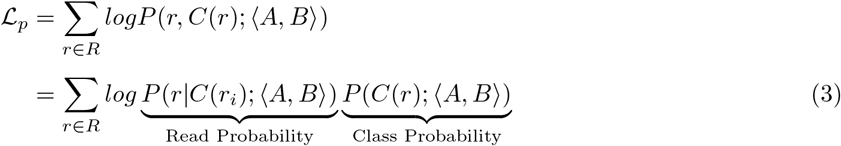

Finally, in a diploid model we assume each read pair is equally likely to originate from allele *A* or *B*:

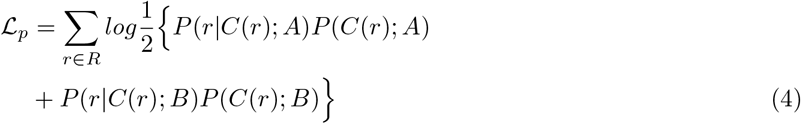

#### 2.2.2. Class probability

The class probability, *P* (*C*(*r*); *A*), models the relative abundance of different classes of informative reads for an underlying repeat length *A*. We use the schematic in Figure 2 to describe how class probabilities are modeled. We consider a repeat with *A* copies of a motif of size *m* bp plus *F* bp of flanking region on either side. Denote the starting position of each read in a pair relative to the beginning of this region as *S*1 and *S*2, where each read in the pair has length *r*. Then we can define class probabilities as:

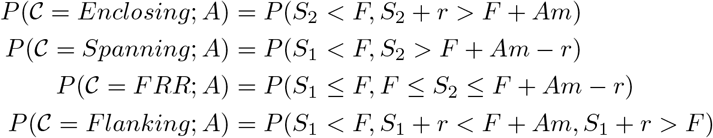

Class probabilities capture changes in the relative abundance of each class as a function of TR length (Supplementary Figure 1). Closed form solutions to compute class probabilities are given in the Supplementary Note.

#### 2.2.3 Read probability

The read probability, *P* (*r|C*(*r*); *A*), models a separate informative characteristic for each class of informative read pairs as a function of repeat length *A*.

The number of repeats observed in enclosing reads (parameter *n* in Figure 2A) can trivially estimate repeat size. However, errors introduced during PCR can alter the number of repeats observed. We model the size of PCR errors using a geometric distribution with default parameter *p* = 0.9 as suggested by HipSTR [44] (Figure 2B).

Spanning read pairs have one mate aligned to either side of the TR. In a sample with a TR expansion, the spanning read pair’s apparent fragment length based on mapped read positions (parameter ∆ in Figure 2C) will shrink compared to the actual fragment length by an amount corresponding to the size of the TR expansion. We thus model observed fragment length as a normal distribution where the mean is a function of repeat length (Figure 2D).

Fully repetitive reads (FRRs) often have an anchor mate that maps in the flanking region before or after the TR. The distance of the anchor from the TR locus (parameter Ω in Figure 2E) is modeled as a function of TR length, with smaller Ω values indicating longer TRs (Figure 2F).

Flanking reads partially cover the TR. The number of repeats in a flanking read (parameter *k* in Figure 2G) indicates that one allele is at least of size *k*. For a TR with length *A*, flanking reads are equally likely to exhibit a number of repeats *k* ranging from 1 to *A* (Figure 2H).

Closed form solutions to compute each class probability are given in the Supplementary Note.

#### 2.2.4 Repetitive Read Count Term

The 𝓛_*N*_ term in (1) assigns a likelihood to the total number of observed fully repetitive reads. We use a Poisson distribution with parameter *λ* to model the expected number of observed FRR reads, which is linearly related to the size of alleles *A* and *B*. Assuming uniform average coverage *C_v_*, read length *r*, and motif length *m*, we can calculate *λ* using (5). The unit step function *u*(.) ensures alleles shorter than the read length have 0 expected FRR reads.

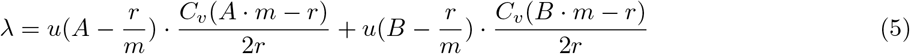

Then we compute the 𝓛_*N*_ term as:

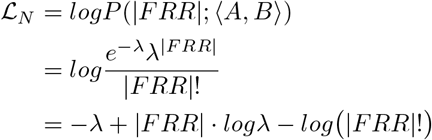

We use Stirling’s approximation to calculate log(*|FRR|*!) for large *|FRR|* values:

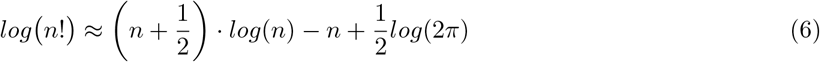

### 2.3 Local Realignment

For enclosing, flanking, and FRR reads GangSTR must obtain accurate counts of the number of repeats contained in each read. For reads fully enclosing the TR plus a minimum of 20bp on either end, repeat count is extracted from the CIGAR score present in the BAM files. Reads starting or ending closer to the TR boundaries or that are fully repetitive are prone to alignment errors and are subject to stringent local realignment. Similar to Tredparse [38], we create artificial reference sequences consisting of flanking region (of size of the read length) on either side and different numbers of repeats, starting from the longest stretch of perfect copies of the repeat and ending with the 2 + 1.1 times the total number of copies of the motif seen in the read. Each read is realigned to the candidate sequences and the reference with the highest realignment score is used to determine the number of repeat copies and the class of the read (flanking, enclosing, or FRR). Realignment is performed using an efficient implementation of the Smith-Waterman algorithm [48].

### 2.4 Retrieving reads mapped to off-target regions

For large expansions some fragments consist entirely of the repeat and may not map to the correct genomic region (off-target). To rescue these reads, we scan a predefined set of off-target regions for additional FRR reads. While in some cases these off-target FRRs cannot be uniquely mapped, our genome-wide analysis below suggests expansions of most TR motifs are rare, and thus most off-target FRRs of the same motif likely originate from the same locus.

To identify off-target regions for each pathogenic TR, we simulated reads for expanded alleles and aligned them back to the reference genome (see simulation settings below). We extracted positions of reads mapped outside of the simulated region (5000bp on either side of the TR). We merged off-target regions within 30bp of each other and expanded the final merged regions by 10bp on either side. The GangSTR implementation allows users to choose whether or not to include off-target FRRs in the maximum likelihood calculation.

### 2.5 Optimization

For each TR, GangSTR determines the possible range of repeat lengths from oberved reads. Minimum and maximum counts are determined by enclosing and flanking reads if present. If FRRs are observed, the maximum count is leniently set to a value with mean expected FRR count 5 times the observed count.

By default, GangSTR uses an exhaustive grid search over all possible allele pairs and returns the maximum likelihood diploid genotype. To speed up optimization for TRs with a large range of possible alleles, GangSTR also implements an efficient multi-step optimization procedure. To account for the irregularity of the likelihood surface, we perform a modular optimization procedure with each step searching a different range of allele lengths. First, any enclosing allele *a* with support of two or more reads is added to the list of potential alleles. In the second step, each potential enclosing allele, *a*, is used to perform 1-dimensional optimization of the likelihood function to find allele *b*, were < *a, b* > minimizes the likelihood function. Next, multiple rounds of 2-dimensional optimization are performed to find < *c, d* > genotypes that minimize the likelihood function. In each round the optimizer uses a different initial point which helps prevent reporting local optima. Any potential allele from each step, *a, b, c, d*, is added to the list of potential alleles. In the final step we compare the likelihood from any combination of two alleles in this list, to find the maximum likelihood genotype. All 1 and 2-dimensional optimization is performed using the COBYLA algorithm [31] implemented in the NLopt library [20].

### 2.6 Quality metrics

GangSTR reports three separate quality metrics to accommodate a range of downstream applications.

#### 2.6.1 Bootstrap confidence intervals and standard errors

In each bootstrap round, GangSTR resamples the set of informative reads (with replacement) to create a bootstrap sample and performs the above optimization procedure on this set of read pairs. The number of bootstrap samples, *N*_*b*_, is set by the user. GangSTR records all bootstrap estimates in separate lists for shorter and longer alleles. These lists are then sorted and used to find the confidence interval at the desired level of significance and standard errors on allele lengths.

#### 2.6.2 Genotype likelihoods and quality score

Let *L* equal the sum of likelihoods for each possible genotype and *L*_*ML*_ be the likelihood of the maximum likelihood genotype. GangSTR returns a quality score 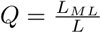. This is equivalent to a posterior probability of the maximum likelihood genotype assuming a uniform prior. This value is most informative for short allele lengths where repeat unit resolution can be achieved. For TR expansions with larger standard errors, the posterior probability of any particular genotype will be low and expansion probabilities are more informative.

#### 2.6.3 Expansion probability

Given a user-specified repeat number expansion threshold *X*, GangSTR computes the probabilities of no expansion (*P*_0_), a heterozygous expansion above the threshold (*P*_1_), or a homozygous expansion above the threshold (*P*_2_) as:

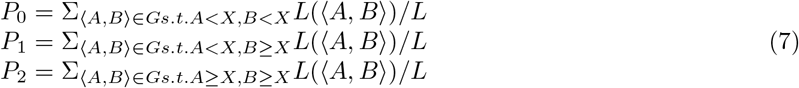

where *G* is the set of all possible diploid genotypes, *L*(〈*A, B*〉) is the likelihood of genotype 〈*A, B*〉, and *L* is as defined above.

### 2.7 Benchmarking using simulated reads

Reads were simulated using wgsim (https://github.com/lh3/wgsim). Unless otherwise specified, we used parameters mean fragment length (-d) 500, standard deviation of fragment length (-s) 100, and read length (−1 and −2) 150. Mutation rate (-r), fraction of indels (-R) and probability of indel extension (-X) were all set to 0, and base error rate (-e) was set to 0.005. The number of simulated reads (-N) was calculated using the following formula 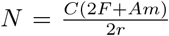, where *C* is the average coverage, set to 40x. *F* is the length of the simulated flanking region around the TR, set to 10,000bp. *A* is the number of copies of the motif of length *m* present in the simulated sample (simulated allele), and *r* is the read length. Simulated genotypes for each pathogenic TR were selected such that the shorter allele covers the normal or premutation range, while the longer allele could be either normal, premutation, or pathogenic (Supplementary Table 1).

Reads were aligned to the hg38 reference genome using BWA-MEM [24] with parameter -M. GangSTR v2.3 was run using the disease-specific reference files for each TR available on the GangSTR website with --coverage set to the simulated coverage level and with the --targeted option. Tredparse v0.7.8 was run with with --cpus 6, --useclippedreads, and --tred appropriately set for each disease locus. ExpansionHunter v2.5.5 was used with and --read-depth preset to the simulated coverage level.

### 2.8 Quantifying genotyping performance with RMSE

Root mean square error (RMSE) was used to compare estimated vs. expected repeat allele lengths. We denote the diploid genotype of sample *i* with 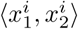. For each diploid genotype, we ordered the two alleles by length such that 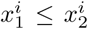. Then to compare estimated 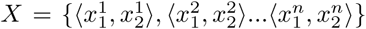 and expected 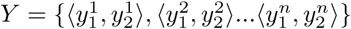 genotypes, RMSE is defined as: 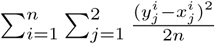

### 2.9 Analysis of genomes and exomes with validated expansions

Whole genome sequencing datasets for samples with previously validated repeat expansions were obtained from the European Genome-Phenome Archive (dataset ID: EGAD00001003562). GangSTR v2.3 was run using the disease-specific reference files for each TR with option --targeted. For Fragile X Syndrome, --ploidy was set to 1 for males and 2 for females. ExpansionHunter v2.5.5 was run using the set of off-target regions given in the GangSTR reference files for Huntington’s Disease, and with their published off-target regions for Fragile X Syndrome. Tredparse v.0.7.8 was run using default parameters and --tred set to HD or FXS for Huntington’s or Fragile X Syndrome, respectively.

Whole exome sequencing datasets for Huntington’s Disease patients were obtained from dbGaP accession phs000371.v1.p1. Fastq files were aligned to the hg19 reference genome using BWA-MEM [24]. PCR duplicates were removed using the samtools [25] rmdup command. Validated repeat lengths were obtained from data fields HDCAG1 and HDCAG2 in table pht002988.v1.p1.c1. We inferred fragment length mean and standard deviation per sample after removing read pairs mapping more than 1kb apart. GangSTR v2.3 was run with --insert-mean and --insert-sdev set to the values computed for each sample. We additionally used parameters --nonuniform and --targeted. ExpansionHunter v2.5.5 was run with --read-depth set to the mean coverage at the TR plus surrounding region. Tredparse v0.7.8 was run with options --useclippedreads and --tred HD.

### 2.10 Constructing a genome-wide repeat reference panel

Tandem Repeats Finder [6] was used to create a panel of repetitive regions with motifs up to 20bp in the hg19 and hg38 reference genomes using parameters matching weight=2, mismatch penalty=5, indel score=17, match probability=80, and indel probability=10. We required a minimum score threshold of 24 to ensure at least 12bp matching the motif for each TR and removed TRs with reference lengths greater than 1000bp.

This initial panel was subject to multiple filters to avoid imperfect or complex TR regions that cannot be accurately genotyped. First, motifs formed by homopolymer runs (i.e., “AAAA”) or by combining smaller sub-motifs (i.e., “ATAT” is made of 2 × “AT”) were discarded. Based on thresholds used in previous TR references [43], we required TRs with motif size 2 or 3 to have at least 5 or 4 copies in tandem, respectively, and larger motifs to have at least 3 copies. To avoid errors in the local realignment step of GangSTR, all repeating regions were trimmed until they no longer contained any imperfections in their first and last three copies of the motif. We removed TRs within 50bp of another TR as these regions tend to be low complexity and result in low quality calls. Next we discarded remaining TRs that do not consist of perfect repetitions of the motif. Finally, we manually added disease associated TRs to ensure notation is consistent with other methods (e.g. [11, 38]).

### 2.11 Run time evaluation

All timing and memory experiments were tested in a Linux environment running Centos 7.4.1708 on a server with 28 cores (Intel^®^ Xeon^®^ CPU E5-2660 v4 @ 2.00GHz) and 125 GB RAM and were performed on a single core. Each experiment was run 5 times and the mean value was reported. Tredparse was evaluated on all available TRs (“treds”) since it does not allow specifying a subset of TRs for analysis. For all timing analyses we used the --skip-unaligned option for ExpansionHunter, which improved run time. For scalability tests, we randomly chose varying sized sets of TRs from the genome-wide reference. Timing was performed with the UNIX time command and the sum of the sys and user times was reported. Memory usage was measured using the UNIX top command. Virtual memory was measured every 0.1 seconds and the maximum value was reported.

### 2.12 Genome-wide TR analysis in a CEU trio

Whole genome sequencing data (BAM files) for the CEU trio consisting of NA12878, NA12891, and NA12892 were obtained from the European Nucleotide Archive (ENA accession: PRJEB3381).

GangSTR v2.3 was run on each family member (NA12878, NA12891, NA12892) using the hg19 ver13 reference available on the GangSTR website with default parameters. We supplied an --str-info-file with the expansion threshold for each TR set to the read length of 101bp. HipSTR v.0.6.2 was run on NA12878 with non-default parameters: --lib-from-samp --def-stutter-model --max-str-len 1200 --min-reads 15 --output-filters. STRetch v0.4.0 was run on NA12878 using the GangSTR reference (limited to motifs up to 6bp) as input regions and with no control genomes specified.

We used our filtering tool, DumpSTR (see Code Availability), to filter GangSTR and HipSTR calls. DumpSTR has various recommended filtering settings depending on the downstream application. For example, for applications where precise estimation of TR length is important, more stringent quality filters should be applied vs. for applications targeted at identifying whether a TR is expanded or not. Thus we applied two filter levels referenced in the results as level 1 and level 2.

First, level 1 filters were used to filter out TRs that could not be reliably called. For HipSTR level 1 filtering, we applied dumpSTR options: --max-call-DP 1000 --min-supp-reads 1, which removes calls with abnormally high coverage or calls with no supporting reads, respectively. For GangSTR level 1 filtering, we applied dumpSTR options: --max-call-DP 1000 --min-call-DP 20 --filter-spanbound-only --filter-badCI, which removes calls with abnormally high coverage, calls where only spanning or bounding reads were found, or calls for which the maximum likelihood genotype falls outside of the 95% bootstrap confidence interval. We additionally filtered regions overlapping annotated segmental duplications in hg19 (UCSC Genome Browser [21] track hg19.genomicSuperDups table).

Second, level 2 filters were used to further restrict to TRs with high confidence length estimates to compare HipSTR vs. GangSTR concordance. For HipSTR level 2 filtering, we applied additional options: --min-call-DP 10 --min-call-Q 0.9 --max-call-flank-indel 0.15 --max-call-stutter 0.15 as recommended on the HipSTR website. For GangSTR level 2 filtering, we applied additional options: –min-call-Q 0.9 –min-total-reads 50.

Mendelian inheritance was determined using two metrics. First, we used maximum likelihood estimates for each sample at each locus to determine whether the child genotype could be explained by parental genotypes. Second, in a less stringent analysis, we determined whether reported confidence intervals were consistent with Mendelian inheritance. Let child, mother, and father confidence intervals be denoted as 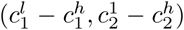, 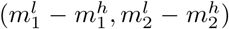, and 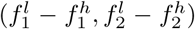, where superscripts 1 and 2 denote the short and long allele at each diploid genotype and subscripts *l* and *h* represent the low and high end of the confidence interval for each allele. A locus was considered to follow Mendelian inheritance if 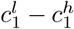 overlapped either maternal confidence interval and 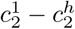 overlapped either paternal confidence interval, or vice versa.

### 2.13 Validating GangSTR using long reads

Oxford Nanopore Technologies (ONT) data for NA12878 was obtained from the Nanopore WGS Consortium (https://github.com/nanopore-wgs-consortium/NA12878). Pacific Biosciences (PacBio) data for NA12878 was obtained from the Genome in a Bottle website (ftp://ftp-trace.ncbi.nlm.nih.gov/giab/ftp/data/NA12878/NA12878_PacBio_MtSinai).

For each repeat, we used the Pysam (https://github.com/pysam-developers/pysam) python wrapper around htslib and samtools [25] to identify overlapping PacBio or ONT reads and extract the portion of the read overlapping the repeat +/-50bp. We estimated the repeat length by taking the difference in length between the reference sequence and the number of bases of each read aligned in that region based on the CIGAR score.

### 2.14 Experimental validation of repeat lengths

Candidate TRs with long alleles identified in NA12878 were PCR amplified using GoTaq (Promega #PRM7123) with primers shown in Supplementary Table 5. PCR products were purified using NucleoSpin^®^ Gel and PCR Clean-up (Macherey-Nagel #740609) and analyzed with capillary electrophoresis using an Agilent 2100 Bio-analyzer and an Agilent DNA 1000 kit (#5067-1504).

### 2.15 Code availability

GangSTR is freely available at https://github.com/gymreklab/GangSTR. The dumpSTR filtering tool is available at https://github.com/gymreklab/STRTools.

## 3 RESULTS

### 3.1 GangSTR outperforms existing TR expansion genotypers

We first evaluated GangSTR’s performance by benchmarking against Tredparse [38] and ExpansionHunter [11], two alternative methods for genotyping repeat expansions. We focused on these methods since they output estimated repeat number at both normal and expanded TRs and do not require a control cohort as input (Table 1). We simulated reads for a set of 14 well-characterized repeats involved in repeat expansion disorders. Since almost all known repeat expansion disorders follow an autosomal dominant inheritance pattern, we simulated individuals heterozygous for one normal range allele and a second allele that varied along the range of normal and pathogenic repeat counts (Supplementary Table 1). In each case, paired-end 150bp reads were simulated to a target of 40-fold coverage, a standard setting for clinical-grade whole genomes. Performance at each locus was measured as the root mean square error (RMSE) between true vs. observed alleles (Methods).

GangSTR genotypes showed the most robust performance compared to other tools across a wide range of repeat lengths, with the smallest RMSE for all TRs tested (Figure 3A, Supplementary Figure 2). At TRs for which the normal range allele is below the read length (SCA6, SCA2, SCA7, SCA1, HTT, and SCA17), both ExpansionHunter and GangSTR accurately predicted the lengths of both alleles (Figure 3B). However, GangSTR demonstrated a distinct advantage over ExpansionHunter in genotyping TRs for which both the normal and pathogenic allele were close to or longer than the read length, where ExpansionHunter estimates become unstable (Figure 3A, C). Tredparse performed well at short alleles but consistently underestimated alleles longer than the fragment length (Figure 3B, C) which accounts for its inflated RMSE results.

**Figure 3:**
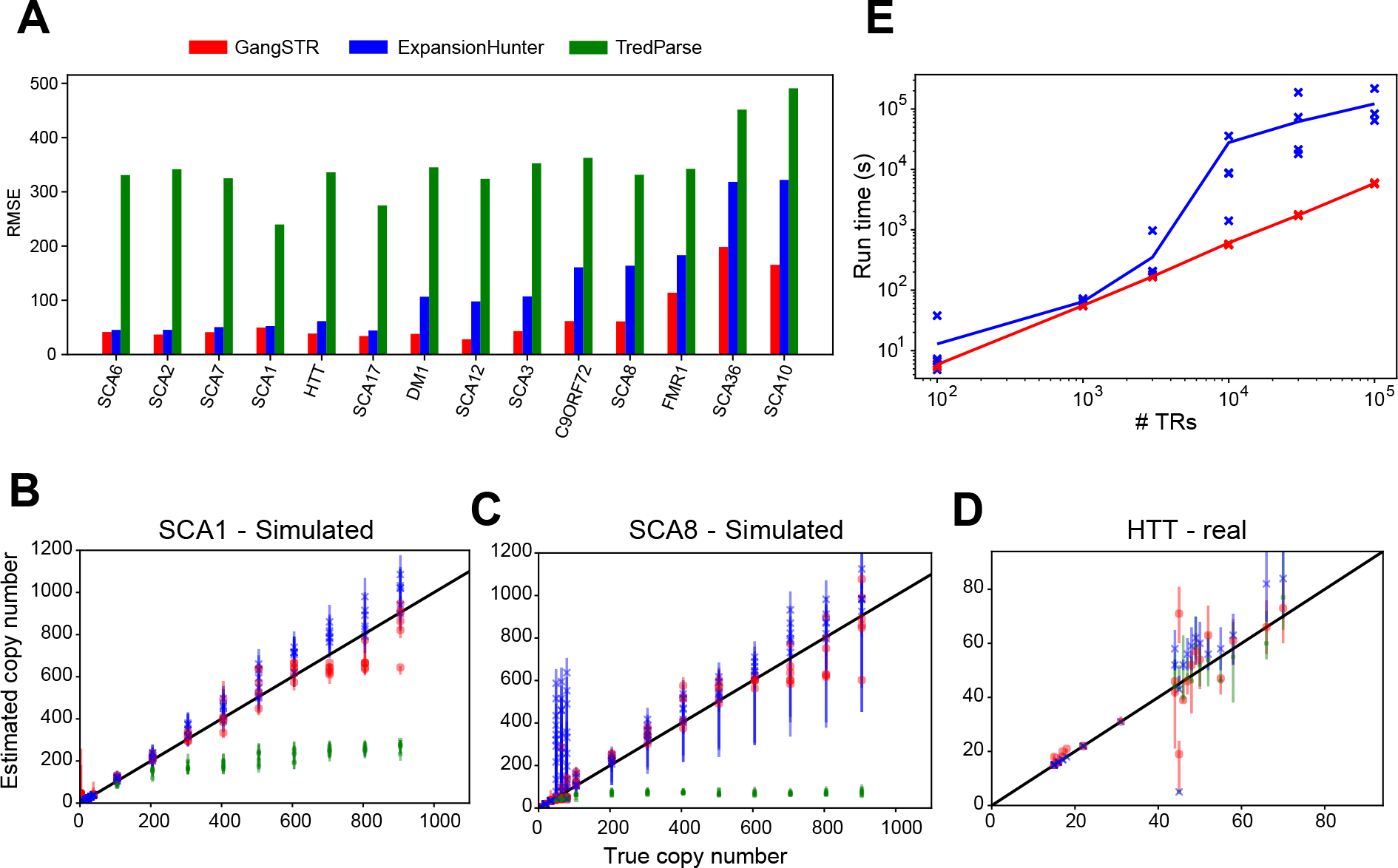
Evaluation of TR genotypers on real and simulated data at pathogenic repeat expansions. A. RMSE for each simulated locus. HTT=Huntington’s Disease; SCA=spinocerebellar ataxia; DM=Myotonic Dystrophy; C9ORF72=amyotrophic lateral sclerosis/frontotemporal dementia. TRs are sorted from left to right by ascending length of the pathogenic allele. B. Comparison of true vs. estimated repeat number for each simulated genotype for HTT. Black solid line gives the diagonal. C. Comparison of true vs. estimated repeat number for each simulated genotype for SCA8. D. Comparison of true vs. estimated repeat number for HTT using real WGS data. E. Run time vs. number of TRs. The x-axis shows the number of TRs in the reference set used. The y-axis shows running time in seconds. Lines give mean value across 5 runs. Points (“x”) give raw data values for each run. For two runs with 10^5^ TRs ExpansionHunter did not run to completion. In all panels, red=GangSTR; blue=ExpansionHunter; green=Tredparse.

We performed additional simulations at the Huntington’s Disease locus to test the effects of sequencing parameters on each tool’s performance. GangSTR and ExpansionHunter both improved significantly as a function of coverage and read length, whereas Tredparse was relatively unaffected (Supplementary Figures 3, 4). Performance of all tools was mostly consistent across mean fragment lengths (Supplementary Figure 5).

We then tested GangSTR’s performance on real NGS data from individuals with validated pathogenic repeat expansions (Methods). Notably, only a small number of such samples are available. Thus tests on real data were limited to two TRs implicated in Huntington’s Disease (HTT) and Fragile X Syndrome (FMR1) with sufficient sample sizes. We first genotyped the HTT and FMR1 loci in 14 and 25 samples respectively with available PCR-free WGS data [11]. All tools performed well on the HTT TR (Figure 3D). GangSTR showed the smallest overall error (RMSE_GANGSTR_=7.9; RMSE_TREDPARSE_=8.3; RMSE_EXPANSIONHUNTER_=10.1) with a small bias in ExpansionHunter for overestimating repeat lengths. Performance was notably worse for all tools at FMR1 (Supplementary Figure 6; RMSE_GANGSTR_=29.3; RMSE_TREDPARSE_=34.8; RMSE_EXPANSIONHUNTER_=27.3). Notably, the FMR1 TR has 100% GC content and very few reads mapping directly to the TR could be identified. This highlights a major challenge in calling GC-rich TRs that are still not sequenced well even with PCR-free protocols.

We additionally tested each tool on 200 whole exome sequencing datasets from patients with validated Huntington’s Disease expansions (Methods, Supplementary Figure 7). GangSTR again showed the smallest error (RMSE_GANGSTR_=5.4; RMSE_TREDPARSE_=96.4; RMSE_EXPANSIONHUNTER_=9.1). Notably, ExpansionHunter gave biased estimates, presumably due to uneven coverage profiles in exomes. Tredparse again underestimated calls for alleles approaching the fragment length (mean=200bp).

Finally, we evaluated computational performance of each tool on various sets of input TRs. We first used the 14 pathogenic TRs to time each tool. GangSTR performed the fastest (mean=15.4s), with ExpansionHunter showing similar run time (mean=16.1s). Tredparse was significantly slower (mean=82.4). We then performed additional evaluation of the scalability of GangSTR and ExpansionHunter by testing on input TR sets ranging from 100 to 100,000 TRs (Figure 3E). GangSTR run time scaled linearly with reference size as expected, whereas ExpansionHunter run time grew super-linearly. Notably, ExpansionHunter only finished on 3 out of 5 runs with 10,000 TRs and would not run to completion on larger TR sets, potentially due to stalling at problematic loci. We additionally tested the maximum memory requirement of each method. GangSTR memory usage stayed relatively constant at under 1GB, whereas ExpansionHunter memory usage grew linearly with the number of TRs in the reference set (Supplementary Figure 8).

### 3.2 Genome-wide TR profiling

We next evaluated GangSTR’s utility for genome-wide TR genotyping. To this end, we used Tandem Repeats Finder [6] to construct a set of all STRs (motif length 2-6bp) and short VNTRs (motif length 7-20bp) in the human reference genome (Methods). In total, we identified 829,231 TRs (780,328 autosomal) in hg19 with a mean length of 15.6bp. Of these, 5,828 are found in coding regions (Figure 4A), most of which have lengths that are multiples of 3bp.

**Figure 4:**
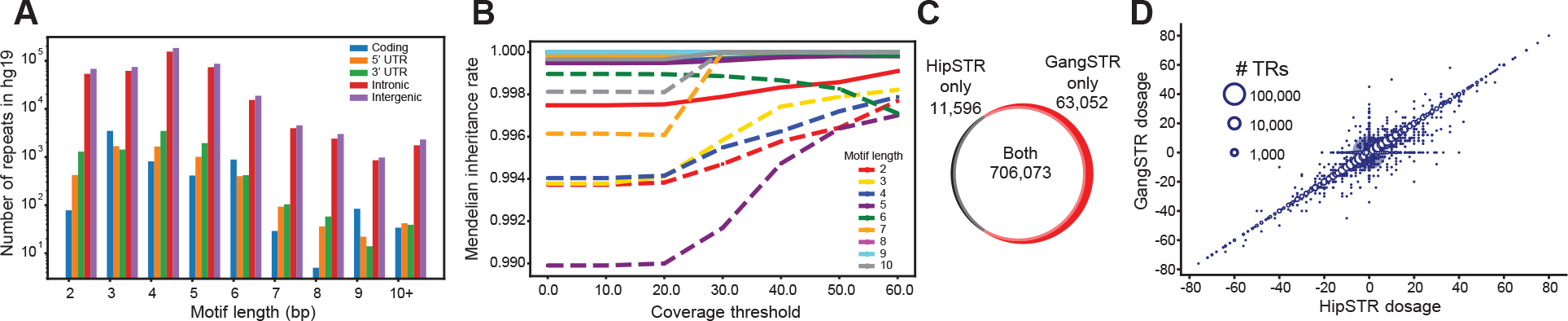
Genome-wide TR genotyping. A. Composition of TRs in the hg19 reference genome. The x-axis gives the motif length and the y-axis (log10 scale) gives the number of TRs in the genome. Colored bars represent TRs overlapping various genomic annotations (blue=coding, orange=5’ UTR, green=3’ UTR, red=intronic, purple=intergenic). B. Mendelian inheritance of GangSTR genotypes in a CEU trio as a function of the number of informative read pairs. Colors denote repeat lengths. Solid lines give mean Mendelian inheritance rate across all TRs, computed based on 95% confidence intervals as described in Methods. Dashed lines are computed after excluding loci where all three samples were homozygous for the reference allele. C. Overlap between TRs genotyped by HipSTR and GangSTR. D. Comparison of HipSTR and GangSTR genotypes. The x-axis and y-axis show the sum of the two allele lengths genotyped by HipSTR and GangSTR in bp relative to the hg19 reference genome (dosage), respectively. The size of the bubble represents the number of points at that coordinate.

We used our genome-wide panel to genotype autosomal repeats using GangSTR on WGS with 30x coverage for a trio of European descent consisting of the highly characterized NA12878 individual and her parents (NA12891 and NA12892). After filtering low quality loci (Methods, level 1 filters), an average of 727,255 TRs were genotyped per sample. As expected, most alleles matched the reference (Supplementary Figure 9) with a bias toward calling alleles shorter than the reference. Both alleles for the majority of TRs (*>*99%) had maximum likelihood lengths less than the read length of 101bp (Supplementary Figure 10). To evaluate GangSTR calls, we determined whether estimated genotypes followed patterns expected based on the trio family structure (Methods). Overall, 98.9% of TRs followed Mendelian inheritance when considering maximum likelihood genotypes. For 99.9% of TRs, 95% confidence intervals were consistent with Mendelian inheritance (Methods). These values changed to 90.6% and 99.3% respectively after removing TRs that were homozygous reference in all samples. The quality of calls steadily increased as a function of the minimum number of observed reads at the locus and was mostly consistent across repeats with different motif lengths (Figure 4B, Supplementary Figure 11).

We evaluated GangSTR’s utility for genome-wide TR profiling by benchmarking against HipSTR using the same reference TR set. After removing low quality loci from each dataset, (Methods, level 1 filters) GangSTR produced calls at 63,052 TRs that could not be reliably genotyped by HipSTR (Figure 4C). The set of TRs not genotyped by HipSTR tended to be longer than average (20.8bp vs. 15.0bp in hg19, onesided t-test *p* < 10^*−*200^) and includes 8/14 known pathogenic TRs analyzed in Figure 3), demonstrating the limitations of relying on enclosing reads. Notably, 11,596 TRs were not called by GangSTR but were present in HipSTR output. These primarily consist of repeats with SNPs or indels in or near the TR sequence which did not pass GangSTR’s stringent local realignment process. After applying stringent recommended quality filters for each tool (Methods, level 2 filters), TRs called by both tools showed extremely high concordance (*>*99%) (Figure 4D) with strong correlation between allele lengths reported by each (Pearson *r*=0.99; *p* < 10^*−*200^; n=651,264), demonstrating that GangSTR can robustly genotype both STRs previously analyzed using HipSTR as well as long TRs previously excluded from genome-wide analyses.

### 3.3 Genome-wide detection of novel TR expansions

We next evaluated whether GangSTR could identify novel repeat expansions in a healthy genome (NA12878). GangSTR identified 69 TRs passing level 1 filters predicted to have at least one allele longer than the read length (101bp) with greater than 80% probability (see Expansion Probabilities defined in Methods) (Supplementary Table 2). Of these, 58 showed evidence of expansions in one or both parents. Long repeats were highly enriched for repeats with motif *AAAG*_*n*_ (27 TRs, one-sided Fisher’s exact test *p* = 2.7 *∗* 10^*−*19^) and related motifs of the form *A*_*n*_*G*_*m*_ (Supplementary Table 3). This finding is concordant with previous reports that *AAG*, *AAAG*, and *AAGG* repeats exhibit strong base-stacking interactions that simultaneously promote expansions through replication slippage and protect the resulting secondary structure from DNA repair [2, 1, 46].

For comparison, we applied STRetch [9], an alternative tool for detecting repeat expansions, using the GangSTR reference TR set of TRs restricted to motif lengths up to 6bp (808,868 total TRs). STRetch leverages a modified reference genome containing decoy repeat sequences to identify potentially expanded TRs. It only attempts to genotype TRs with candidate expansions and thus is unsuitable for unbiased genome-wide TR genotyping. After filtering for segmental duplications (Methods), STRetch returned results for 45 TRs (Supplementary Table 4). Notably, STRetch took approximately 157 CPU-hours (6.5 days) compared to 16.6 CPU-hours for GangSTR on a single genome. TRs genotyped by both GangSTR and STRetch showed concordant repeat number estimates (Pearson *r*=0.61, *p* = 4.9 * 10^*−*5^, n=37, Figure 5A, Supplementary Table 4). However only 5 of the 69 TRs with alleles longer than 101bp reported by GangSTR were genotyped by STRetch. Overall these results show that GangSTR provides a more comprehensive analysis of genome-wide TR variation.

**Figure 5:**
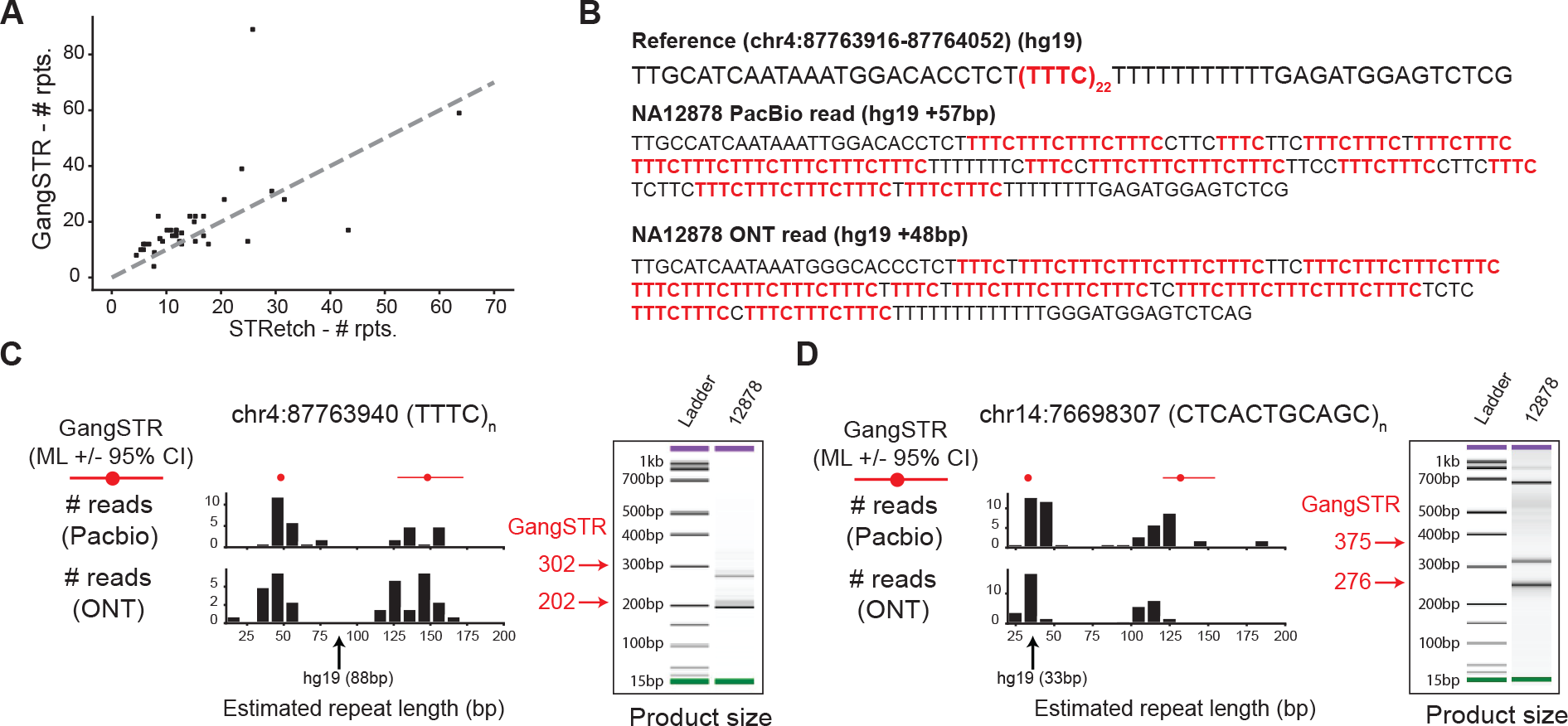
Discovery and validation of genome-wide TR expansions. A. Comparison of STRetch and GangSTR estimated repeat lengths. The x-axis shows the estimated repeat number returned by STRetch. The y-axis shows the estimated repeat number of the longest of two alleles reported as the maximum likelihood genotype by GangSTR. Only TRs called by both tools and passing all GangSTR filters are shown. The gray dashed line shows the diagonal. B. Example sequence at a candidate TR expansion. The reference sequence and representative reads from PacBio (top) and ONT (bottom) for NA12878 are shown for a locus where GangSTR predicted a 48bp expansion from the reference genome. Instances of the repeat motif are shown in red. C, D. For each of the TRs shown, left plots compare GangSTR genotypes to those predicted by long reads. Red dots give the maximum likelihood repeat lengths predicted by GangSTR and red lines give the 95% confidence intervals for each allele. Black histograms give the distribution of repeat lengths supported by PacBio (top) and ONT (bottom) reads. The black arrow denotes the length in hg19. The right plots show PCR product sizes for each TR as estimated using capillary electrophoresis. Left bands show the ladder and right bands show product sizes in NA12878. Green and purple bands show the lower and upper limits of the ladder, respectively. Red arrows and numbers give product sizes expected for the two alleles called by GangSTR.

To validate putative expansions identified by GangSTR, we examined long read data from WGS for NA12878 generated using Pacific Biosciences (PacBio) [28] and Oxford Nanopore Technologies (ONT) [19]. For each of the 69 TRs with at least one allele longer than the read length, we extracted regions of PacBio and ONT reads overlapping the TR and determined the repeat length supported by each read (Methods). In 60/68 cases with supporting reads from PacBio, at least one read showed evidence of an allele *>*101bp (51/68 for ONT) (Supplementary Table 2). ONT showed less evidence of expansions, perhaps due to a deletion bias. Both long read technologies exhibit high error rates at homopolymer runs [42], resulting in messy sequence within repeats themselves (Figure 5B)

Finally, for a subset of 12 candidate expansions, we additionally performed capillary electrophoresis to measure TR lengths (Methods, Supplementary Table 5). Capillary results showed evidence of long alleles for the majority (10/12) of TRs (Figure 5C,D, Supplementary Figure 12). Notably, expanded TRs proved difficult to amplify and capillary results in many cases did not clearly indicate two distinct allele lengths. Further, in some cases GangSTR, PacBio, and ONT gave discordant results, with either strikingly different repeat lengths or an ambiguous signal that could not be resolved using capillary electrophoresis (Supplementary Figure 12). Still, the majority of long TRs identified by GangSTR were validated by at least one of these orthogonal technologies. Taken together, these results demonstrate GangSTR’s ability to identify novel expanded TRs from genome-wide data and highlight the challenges in precisely validating TR lengths at these loci.

## 4 DISCUSSION

### 4.1 A unified framework for genotyping a wide range of TRs

Our study presents GangSTR, a novel tool for genotyping TRs from NGS data. GangSTR is a flexible tool that can be used for a variety of applications, including genome-wide TR genotyping, targeted detection of TR expansions at known pathogenic loci, and genome-wide discovery of novel TR expansions. We show that GangSTR outperforms existing tools in both speed and accuracy in a range of settings using simulated and real NGS datasets. We applied GangSTR genome-wide to genotype hundreds of thousands of TRs in a deeply sequenced healthy trio. We identified dozens of long repeat alleles which were confirmed by orthogonal long read and capillary electrophoresis technologies.

GangSTR outperforms state of the art methods for characterizing TR expansions from NGS (Figure 3). Our targeted simulation analyses demonstrate that GangSTR produced accurate TR length estimates in a range of settings, including unexpanded genotypes and genotypes that are either heterozygous or homozygous for long alleles. GangSTR’s advantage becomes more pronounced for TRs with longer normal-length alleles. ExpansionHunter [11] does not accurately genotype TRs heterozygous for two long alleles since its model is primarily based on sequencing coverage. Our model overcomes this limitation by incorporating orthogonal information available from spanning read pairs. While Tredparse [38] similarly models observed fragment lengths for spanning read pairs, it does not analyze read pairs where both reads are mapped to off-target regions, and cannot genotype TRs longer than the fragment length.

Beyond TR expansions implicated in Mendelian disorders, mounting evidence suggests that thousands of TRs genome-wide contribute to polygenic phenotypes such as gene expression [13]. Accurate genome-wide TR genotyping will be critical for performing association studies to identify these TRs and quantify their contribution to common disease. GangSTR extends our existing methods for genome-wide TR genotyping to accommodate repeats longer than the read length and identifies tens of thousands of TRs that were missed by HipSTR [44].

Genome-wide analysis additionally allows for identifying novel pathogenic TR expansions or expansions present in healthy genomes. While existing tools allow for this, they do not produce genome-wide TR length estimates. STRetch [10] identifies novel expansions, but requires a time-consuming and memory intensive step to realign raw reads to a modified reference sequence containing decoy regions. Due to compute requirements. performing realignment is often not feasible to implement in high-throughput pipelines. Additionally, STRetch only identifies a subset of TR alleles that are expanded from the repeat sequence, and thus cannot be used to obtain accurate diploid TR lengths. exSTRa [39] can also be used to find novel expansions, but requires a matched control cohort to identify expansions and reports only expansion status, rather than TR length estimates. On the other hand, GangSTR generates unbiased TR length estimates genome-wide, which can be used in diverse downstream applications such as association testing or discovery of Mendelian disease loci. Further, GangSTR is far more efficient, taking around 16.5 CPU-hours to run on a single genome compared to days for competing methods.

### 4.2 Remaining challenges in TR genotyping

Genome-wide TR genotyping still faces several important limitations. First, all tools described here, including GangSTR, require a TR panel based on the reference genome as input. Thus they are not able to genotype TRs that are not properly assembled in the reference genome. Additionally, TRs with complex structures, such as sequence imperfections, highly repetitive flanking regions, or multiple different adjacent repeating motifs, are ambiguous to define and their boundaries depend highly on the choice of parameters used to create the reference. Complex TRs are a source of errors in GangSTR genotypes. Our realignment step relies on aligning reads to an artificial reference created for each possible TR allele by stitching together perfect repetitions of the TR motif. Because of this design choice, repetitive motifs in the flanking regions surrounding a TR locus can reduce robustness of the realignment step. We attempt to filter most of these regions from our reference set to avaoid TRs that cannot be reliably called. A more complex model is required to account for these regions.

Most previous tools focused on STRs with motifs up to 6bp. Here, we have expanded our reference to include VNTRs with motifs up to 20bp. This limit can theoretically be expanded. However, longer motifs tend to have more complex imperfections. Additionally, several aspects of GangSTR’s model rely on identifying several copies of a repeat unit in a single read (e.g. enclosing and flanking reads). Thus accuracy is likely to decrease slightly at longer motifs.

Second, due to a lack of large ground truth datasets our validation experiments relied heavily on simulated data. These simulations assume uniform coverage and do not capture many error modes present in real data such as PCR, GC biases, or DNA degradation.

Third, some TRs are still not adequately captured by short reads. For example, TRs in regions with extremely high GC content are often very poorly covered due to biases induced by PCR and other sequencing steps. Furthermore, TRs with highly repetitive flanking regions are still inaccessible due to poor sequence alignment of anchoring or spanning reads. Additionally, while GangSTR can genotype TRs well beyond the fragment length, it still produces noisy estimates at extremely long TRs (e.g. thousands of bp), especially when both alleles are long. We suspect this is primarily due to variance in FRR coverage which grows linearly with total repeat length. While some of these challenges may be overcome with improved modeling techniques, some TRs are likely to remain out of reach using NGS.

Finally, for some repeats we could not obtain reliable genotypes using any technology, including short reads, long reads, or PCR methods. This may be due to a combination of difficulty amplifying highly repetitive regions, difficulty sequencing complex repeats, or high error rates in long read data. Additionally, some unstable repeats may exhibit high rates of somatic variation [37, 22], rendering the notion of a “correct” genotype meaningless. Indeed, for several loci we saw evidence of a spectrum of repeat numbers in all technologies tested. GangSTR could be extended in the future to incorporate somatic mosaicism into its model.

Some of the limitations mentioned above could be overcome using long read technologies such as PacBio or ONT. However, we focused on Illumina short reads here as Illumina is rapidly becoming the clinical standard and remains unmatched in cost and accuracy. It is likely that hybrid approaches combining both short and long read data will provide the greatest accuracy.

## 5 CONCLUSION

Overall, GangSTR allows rapid and accurate genotyping of both short and expanded TRs and can be readily applied to large NGS cohorts to enable novel genetic discoveries across a broad range of applications.

## 6 ACKNOWLEDGEMENTS

We thank Vineet Bafna, Vikas Bansal, Egor Dolzhenko, and Alon Goren for helpful comments on the method and manuscript. We thank Harriet Dashnow for advice on running STRetch.

Whole exome sequencing for Huntington’s Disease patients was obtained from DbGaP accession phs000371.v1.p1, which was supported by the Massachusetts Huntington’s Disease Center without Walls (NS16367) and the PREDICT-HD study of the Huntington Study Group (NS40068).

