## Supplementary Note for "Profiling the genome-wide landscape of tandem repeat expansions"

### Supplementary Material - Mousavi, *et al.*

### 1 Supplementary Note: GangSTR Model

#### 1.1 Class probabilities

Class probability describes the probability of a read pair belonging to a specific class, considering uniform coverage. For any value of underlying allele length $A$, this probability can give an intuition for the relative abundance of different classes of reads **(Supplementary Figure 1)**.

Derivation of class probabilities for each read pair class are given below. Notation corresponds to that used in the main text and depicted in **Figure 2**.

##### 1.1.1 Class Probability of Enclosing Reads

Without loss of generality, we assume the first mate in the pair is enclosing. The calculation is similar for the other mate (Equation (1)). Assuming uniformity of coverage, we use a uniform distribution to find the probability of a TR region being enclosed by a read.

$$\begin{aligned} P(c_i = E; A) &= P(S_1 < F, S_1 + r > F + A \cdot m) \\ &= P(F + A \cdot m - r < S_1 < F) \\ &= \frac{(F) - (F + A \cdot m - r)}{2F + A \cdot m - 2r} \\ &= \frac{(r - A \cdot m)}{2F + A \cdot m - 2r} \end{aligned} \tag{1}$$

##### 1.1.2 Class Probability of Spanning Reads

##### 1.1.3 Fragment Length Distribution

We model the observed fragment length $\delta$ with a limited Gaussian random variable $\Delta$ with the following distribution:

$$f_\Delta(\delta) = \frac{1}{C\sqrt{2\pi}\sigma} e^{-\frac{1}{2\sigma^2}(\delta-\mu)^2} \qquad ; r \leq \delta \leq \infty \tag{2}$$

In this equation $\mu$ is average fragment length, $\sigma$ is the standard deviation of the fragment length distribution, $C$ is a normalization constant to account for limited range of $\delta$, and $r$ is the read length.

Integration of this probability density function arises several times throughout the rest of this document. We compute these integrals using a helper Gaussian distribution X:

$$f_X(x) = \frac{1}{\sqrt{2\pi}\sigma} e^{-\frac{1}{2\sigma^2}(x-\mu)^2} \qquad ; -\infty \leq x \leq \infty \tag{3}$$

and it's cumulative density function (CDF):

$$F_X(x) = \int_{-\infty}^{x} f_X(x)dx \tag{4}$$

1) For $r \leq a, b \leq \infty$:

$$\begin{aligned} \int_a^b f_\Delta(\delta)d\delta &= F_\Delta(b) - F_\Delta(a) \\ &= \frac{1}{C}\{F_X(b) - F_X(a)\} \end{aligned}$$

2) For $r \leq a, b \leq \infty$:

$$\begin{aligned}\int_a^b (\delta-\mu) f_\Delta(\delta) d\delta &= \frac{1}{C\sqrt{2\pi}\sigma}\int_a^b (\delta-\mu) e^{-\frac{1}{2\sigma^2}(\delta-\mu)^2} d(\delta-\mu)\\ &= -\frac{\sigma}{C\sqrt{2\pi}}\{e^{-\frac{1}{2\sigma^2}(b-\mu)^2} - e^{-\frac{1}{2\sigma^2}(a-\mu)^2}\}\\ &= -\frac{\sigma^2}{C}\{f_X(b) - f_X(a)\}\end{aligned}$$

A read pair is classified as spanning if it's two mates are mapped in the flanking region before and after the TR locus.

$$\begin{aligned}P(c_i = S; A) &= P(S_1 < F, S_2 > F + A\cdot m - r)\\ &= \int_{2r}^{2F+A\cdot m} P(S_1 < F, S_2 > F + A\cdot m - r|\Delta = \delta) f_\Delta(\delta) d\delta\\ &= \int_{2r}^{2F+A\cdot m} P(S_1 < F, S_1 + \Delta - r > F + A\cdot m - r|\Delta = \delta) f_\Delta(\delta) d\delta\\ &= \int_{2r}^{2F+A\cdot m} P(F + A\cdot m - \Delta < S_1 < F)|\Delta = \delta) f_\Delta(\delta) d\delta \qquad (5)\\ &= \int_{2r}^{2F+A\cdot m} \frac{(F) - (F + A\cdot m - \delta)}{2F + A\cdot m - 2r} u(\delta - A\cdot m) f_\Delta(\delta) d\delta \qquad (6)\\ &= \int_{max\{2r, A\cdot m\}}^{2F+A\cdot m} \frac{\delta - A\cdot m}{2F + A\cdot m - 2r} f_\Delta(\delta) d\delta\end{aligned}$$

Step function $u(.)$ is introduced in (6) to satisfy the condition in (5), $x + A\cdot m - \Delta < x$, which simplifies to $\Delta > A\cdot m$. This condition is then imposed in the integral limit. We continue the calculation using the helper integrals from Section 1.1.3.

$$\begin{aligned}P(c_i = S; A) &= \int_{max\{2r, A\cdot m\}}^{2F+A\cdot m} \frac{(\delta-\mu) + \mu - A\cdot m}{2F + A\cdot m - 2r} f_\Delta(\delta) d\delta\\ &= \frac{1}{2F + A\cdot m - 2r}\Big\{(\mu - A\cdot m)\int_{max\{2r, A\cdot m\}}^{2F+A\cdot m} f_\Delta(\delta) d\delta\\ &+ \int_{max\{2r, A\cdot m\}}^{2F+A\cdot m} (\delta - \mu) f_\Delta(\delta) d\delta\Big\}\\ &= \frac{\mu - A\cdot m}{C(2F + A\cdot m - 2r)}\Big[F_X\big(2F + A\cdot m\big) - F_X\big(max\{2r, A\cdot m\}\big)\Big]\\ &- \frac{\sigma^2}{C(2F + A\cdot m - 2r)}\Big[f_X\big(2F + A\cdot m\big) - f_X\big(max\{2r, A\cdot m\}\big)\Big]\\ &= \frac{1}{C(2F + A\cdot m - 2r)}\Bigg\{(\mu - A\cdot m)\Big[F_X\big(2F + A\cdot m\big) - F_X\big(max\{2r, A\cdot m\}\big)\Big]\\ &- \sigma^2\Big[f_X\big(2F + A\cdot m\big) - f_X\big(max\{2r, A\cdot m\}\big)\Big]\Bigg\}\end{aligned}$$

##### 1.1.4 Class Probability of Flanking Reads

Without loss of generality, we assume the first mate in the pair is flanking. The calculation is similar for the other mate. Assuming uniform coverage, we use a uniform distribution to find the probability of observing a flanking read.

$$
\begin{aligned}
P(c_i = F; A) &= P\big(S_1 < F, S_1 + r < F + A \cdot m, S_1 + r > F\big) \\
&= P\big(F - r < S_1 < min\{F, F + A \cdot m - r\}\big) \\
&= \frac{min\{F + A \cdot m - r, F\} - (F - r)}{2F + A \cdot m - 2r} \\
&= \frac{F + min\{A \cdot m - r, 0\} - F + r}{2F + A \cdot m - 2r} \\
&= \frac{min\{A \cdot m, r\}}{2F + A \cdot m - 2r}
\end{aligned} \tag{7}
$$

##### 1.1.5 Class Probability of FRRs

$$
\begin{aligned}
P(c_i = FRR; A) &= P(S_1 \leq F, F \leq S_2 \leq F + A \cdot m - r) \\
&= \int_{2r}^{2F + A \cdot m} P(S_1 \leq F, F \leq S_1 + \Delta - r \leq F + A \cdot m - r | \Delta = \delta) f_\Delta(\delta) d\delta \\
&= \int_{2r}^{2F + A \cdot m} P(S_1 \leq x, S_1 \leq F + A \cdot m - \delta, S_1 \geq x + r - \delta) f_\Delta(\delta) d\delta
\end{aligned} \tag{8}
$$

We combine the inequalities describing $S_1$ in (8) to derive conditions that need to hold for this integral to have non-zero value.

- $\left.\begin{matrix} S_1 \geq F + r - \delta \\ S_1 \leq F + A \cdot m - \delta \end{matrix}\right\} \Rightarrow F + A \cdot m - \delta \geq F + r - \delta \Rightarrow A \cdot m \geq r$

  $\Rightarrow$ This condition is the clear condition underlying presence of FRR reads. Smaller TR lengths have 0 probability of having an FRR read.

- $\left.\begin{matrix} S_1 \geq F + r - \delta \\ S_1 \leq F \end{matrix}\right\} \Rightarrow F \geq F + r - \delta \Rightarrow \delta \geq r$

  $\Rightarrow$ The lower limit of the integral is $\delta \geq 2r$, hence this condition is satisfied for the range of possible $\delta$ values.

Since there are two upper bounds for $S_1$ in (8), we need to consider two different scenarios:

- $x \leq F + A \cdot m - \delta \Rightarrow \delta \leq A \cdot m$
  Therefore, for $2r \leq \delta \leq A \cdot m$ ; $A \cdot m \geq 2r$, integrand is simplified to:
  $\Rightarrow P(S_1 \leq F, S_1 \leq F + A \cdot m - \delta, S_1 \geq F + r - \delta) = P(F + r - \delta \leq S_1 \leq F)$
  For $A \cdot m < 2r$, this part has no contribution.

- $F > F + A \cdot m - \delta \Rightarrow \delta > A \cdot m$
  Similarly, for $A \cdot m \leq \delta \leq 2F + A \cdot m$, integrand is simplified to:
  $\Rightarrow P(S_1 \leq F, S_1 \leq F + A \cdot m - \delta, S_1 \geq F + r - \delta) = P(F + r - \delta \leq S_1 \leq F + A \cdot m - \delta)$

Continuing integration for $A \cdot m \geq 2r$:

$$\begin{aligned}
P(c_i = FRR; A) &= \int_{2r}^{A\cdot m} P(F + r - \delta \leq S_1 \leq F) f_\Delta(\delta) d\delta \\
&+ \int_{A\cdot m}^{2F+A\cdot m} P(F + r - \delta \leq S_1 \leq F + A \cdot m - \delta) f_\Delta(\delta) d\delta \\
&= \int_{2r}^{A\cdot m} \frac{(F) - (F + r - \delta)}{2F + A \cdot m - 2r} f_\Delta(\delta) d\delta \\
&+ \int_{A\cdot m}^{2F+A\cdot m} \frac{(F + A \cdot m - \delta) - (F + r - \delta)}{2F + A \cdot m - 2r} f_\Delta(\delta) d\delta \\
&= \int_{2r}^{A\cdot m} \frac{(\delta - \mu) + (\mu - r)}{2F + A \cdot m - 2r} f_\Delta(\delta) d\delta \\
&+ \int_{A\cdot m}^{2F+A\cdot m} \frac{A \cdot m - r}{2F + A \cdot m - 2r} f_\Delta(\delta) d\delta \\
&= \frac{1}{C(2F + A \cdot m - 2r)} \Big\{ -\sigma^2 \big[f_X(A \cdot m) - f_X(2r)\big] \\
&+ (\mu - r)\big[F_X(A \cdot m) - F_X(2r)\big] \\
&+ (A \cdot m - r)\big[F_X(2F + A \cdot m) - F_X(A \cdot m)\big] \Big\} \qquad ; A \cdot m \geq 2r
\end{aligned}$$

The result is similar for $A \cdot m < 2r$, except the first two terms are zero in this case:

$$P(c_i = FRR; A) = \frac{A \cdot m - r}{C(2F + A \cdot m - 2r)} \Big\{ F_X(2F + A \cdot m) - F_X(A \cdot m) \Big\} \quad ; A \cdot m < 2r \tag{9}$$

#### 1.2 Read probabilities

For each class of informative reads, the read probability describes the distribution of the informative characteristic of the class, given an underlying allele $A$ **(Figure 2)**. The details of read probability for each class of informative reads is presented in the following sections.

##### 1.2.1 Enclosing Reads

Enclosing reads contain the whole repeating region, as well as flanking regions before and after. Therefore, the number of copies can be directly extracted after performing the local realignment step..

The HipSTR stutter model [1] explains the distribution of the number of repeat copies in enclosing reads. Equation (10) shows the probability of a read with $r_i$ copies having an error of length $\delta$ copies compared to the underlying true number of copies $A$. In this model, $u$ and $d$ correspond to the probability of stutter adding or removing copies of the motif, and $\rho_s$ is the parameter of the geometric distribution that governs the number of stutter deviations from true number of copies $A$.

$$P(r_i - A = \delta | c_i = E; A) = \begin{cases} 1 - u - d & \delta = 0 \\ u\rho_s (1 - \rho_s)^{\delta - 1} & \delta > 0 \\ d\rho_s (1 - \rho_s)^{-\delta - 1} & \delta < 0 \end{cases} \tag{10}$$

##### 1.2.2 Flanking Reads

Flanking reads with $n$ copies of the motif imply that one of the alleles has at least $n$ copies of the motif. We use a uniform distribution (similar to [2]) to model the distribution of reads in the flanking class:

$$P(r_i = n | c_i = F; A) = \begin{cases} \frac{1}{A} & n \leq A \\ 0 & n > A \end{cases} \tag{11}$$

##### 1.2.3 Spanning Reads

Fragments that completely span the TR region can create spanning read pairs. Spanning read pairs consist of two mates that are mapped to the flanking region before and after the TR. During alignment, spanning reads originating from an expanded TR allele experience a decrease in the observed fragment length (**Figure 2C-D**). Therefore, the distribution of fragment lengths for spanning reads is similar to the fragment length distribution in section 1.1.3, with a decrease in average fragment length by an amount equal to the size of expansion. If the reference has $R$ copies of an $m$ base pair motif, we can describe the class probability of spanning reads with the following Gaussian distribution:

$$P(r_i | c_i = S) \sim N(\mu - (A - R) \cdot m, \sigma) \tag{12}$$

##### 1.2.4 Fully Repetitive Reads (FRRs)

FRR reads are extracted from both on and off target regions to create the repetitive read count term in the likelihood model. Here we discuss another informative aspect of FRR reads, the distance of an anchored mate to the repeat region (**Figure 2**).

Anchored FRRs are read pairs that contain one read completely consisting of repeats, while the other mate pair is mapped to the flanking region before or after the TR. Using the fragment length distribution (see 1.1.3), we model the distance of the anchor read from the repeat region (shown by $\Omega$) to obtain read probability of this class of reads. We use the notation from section 1.1.3 to derive the read probability of anchored FRR reads.

$$\begin{aligned} P(r_i | c_i = FRR; A) &= P(\Omega + 2r < \Delta < \Omega + r + L) && (13) \\ &= F_\Delta(\Omega + r + L) - F_\Delta(\Omega + 2r) && (14) \\ &= \frac{1}{C}[F_X(\Omega + r + L) - F_X(\Omega + 2r)] && (15) \end{aligned}$$

On the other hand, fragments that originate from within the repeating region generate FRR read pairs (both mates repetitive). These read pairs do not have an anchor, and are most likely aligned to one of the off-target regions associated with the TR. These read pairs contribute to both FRR count term (adding two FRR reads) and read pair term (FRR class probability computed for $\Omega = -r$).

### 2 Supplementary Figures

#### Supplementary Figure 1

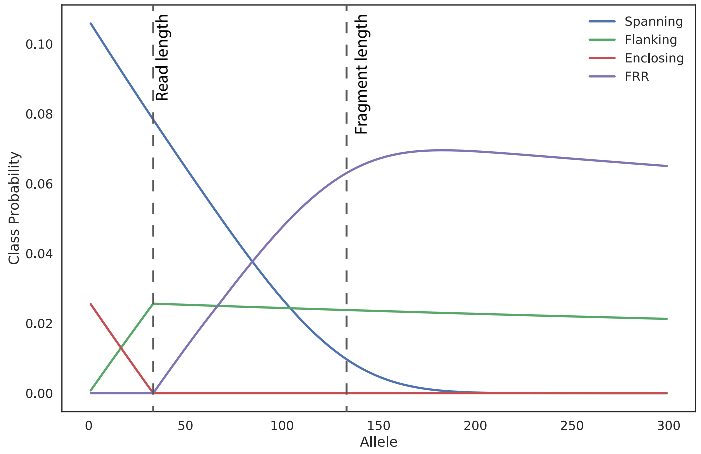


**Class probabilities as a function of TR length.** The x-axis shows the allele length in number of repeats. The y-axis shows the probability that a read mapped to the TR region would be from each class. Results were calculated using a 3bp long repeat unit, read length=100bp, and fragment length=400bp. Blue=spanning reads, green=flanking reads, red=enclosing reads, and purple=FRR reads.

### Supplementary Figure 2

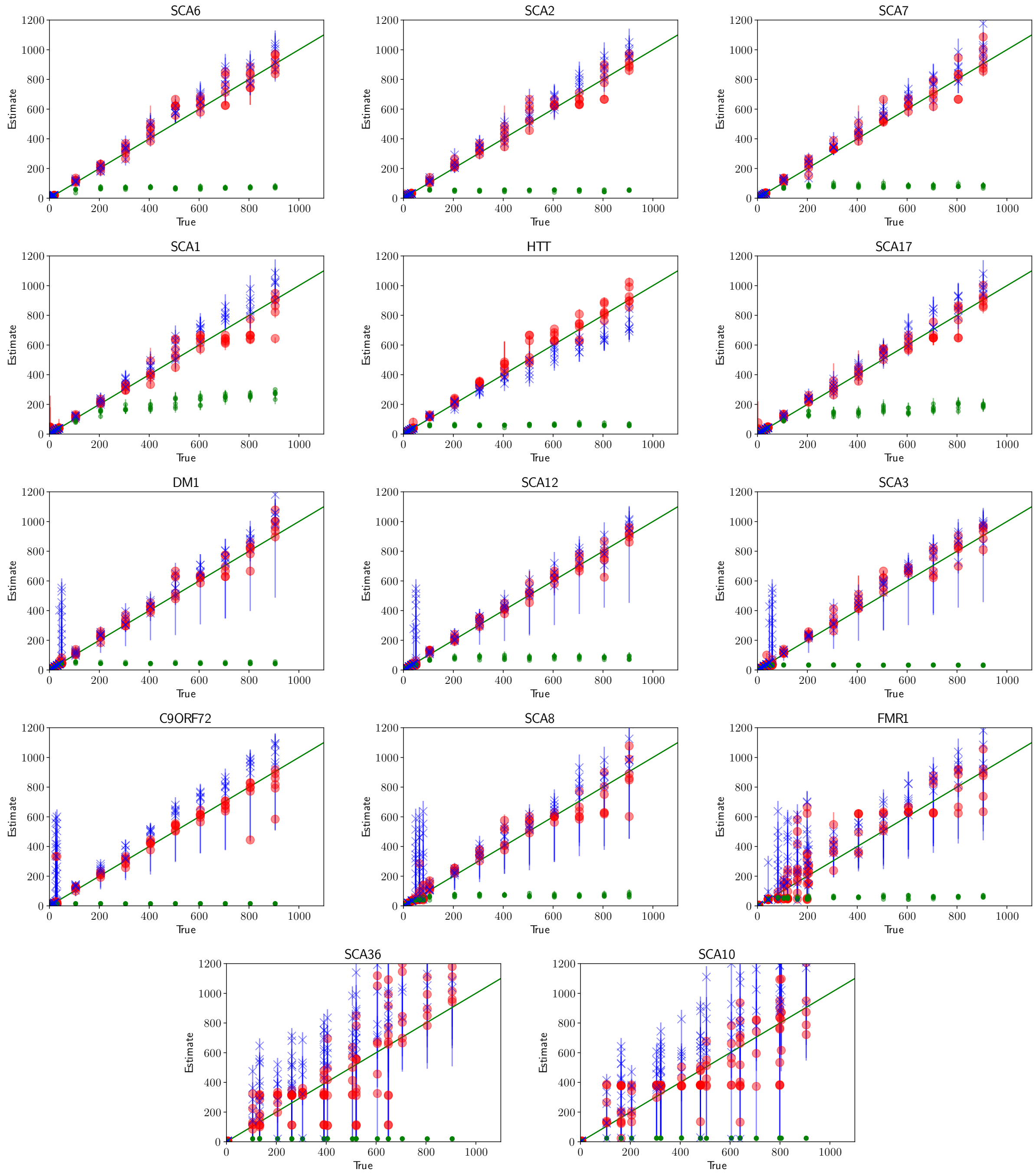


**Comparison of true vs. estimated repeat number on simulated data for different loci** The x-axis shows the simulated allele length in number of repeats. The y-axis shows the estimated allele length in number of repeats. The accuracy (root mean square error) for each panel is plotted in Figure 3A. In all panels, red=GangSTR; blue=ExpansionHunter; green=Tredparse.

#### Supplementary Figure 3

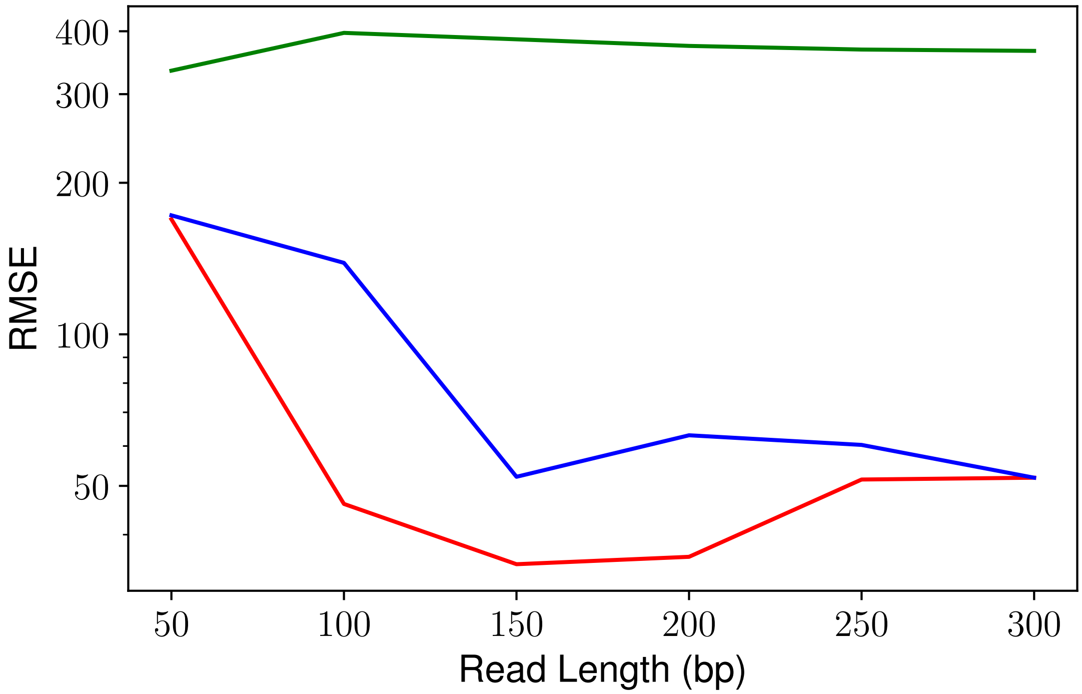


**Estimation accuracy for simulated samples of HTT vs. read length** Root Mean Square Error (RMSE) for estimation of simulated samples of the HTT locus with different read lengths. Red=GangSTR; blue=ExpansionHunter; green=Tredparse.

#### Supplementary Figure 4

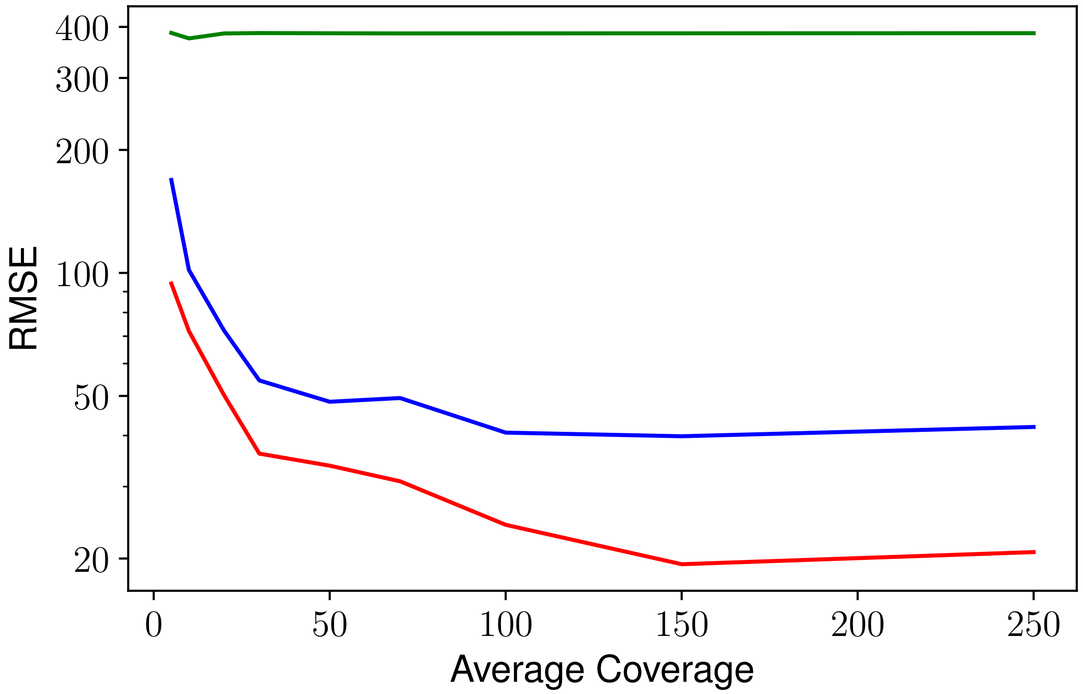


**Estimation accuracy for simulated samples of HTT vs. average coverage** Root Mean Square (RMSE) for estimation of simulated samples of the HTT locus with different coverages. Red=GangSTR; blue=ExpansionHunter; green=Tredparse.

#### Supplementary Figure 5

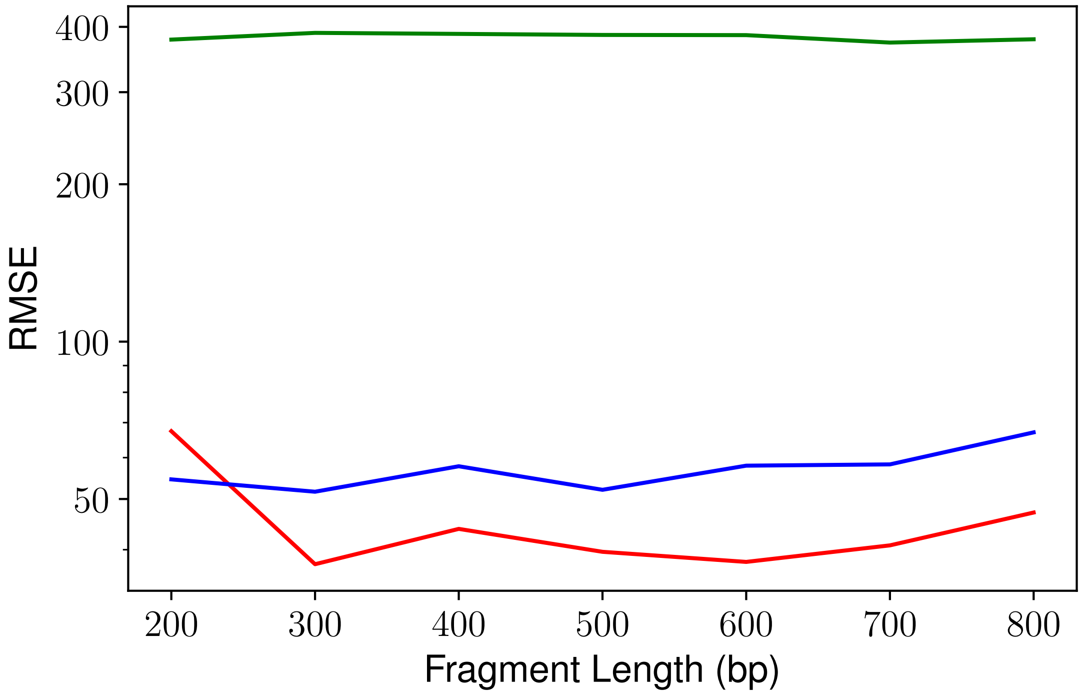


**Estimation accuracy for simulated samples of HTT vs. fragment length** Root Mean Square (RMSE) for estimation of simulated samples of HTT locus with different fragment lengths. Red=GangSTR; blue=ExpansionHunter; green=Tredparse.

### Supplementary Figure 6

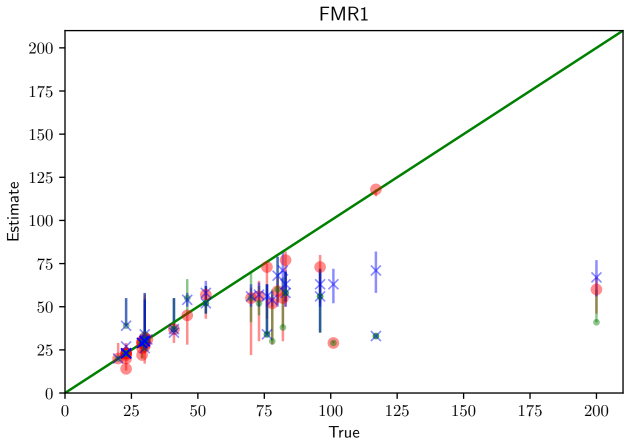


**Comparison of repeat expansion tools at FMR1 using real WGS data.** The x-axis shows the experimentally validated allele length in number of repeats. The y-axis shows the estimated allele length in number of repeats. Green solid line gives the diagonal. red=GangSTR; blue=ExpansionHunter; green=Tredparse.

### Supplementary Figure 7

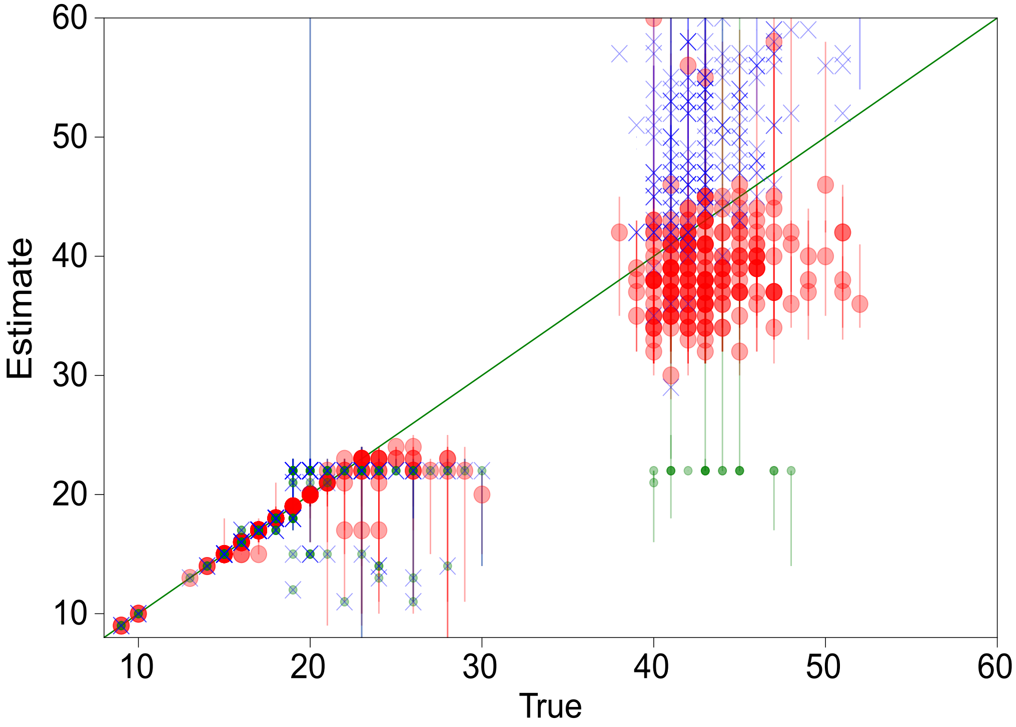


**Comparison of true vs. estimated repeat number using real HTT exome data** The x-axis shows the experimentally validated allele length in number of repeats. The y-axis shows the estimated allele length in number of repeats. Green solid line gives the diagonal. red=GangSTR; blue=ExpansionHunter; green=Tredparse.

### Supplementary Figure 8

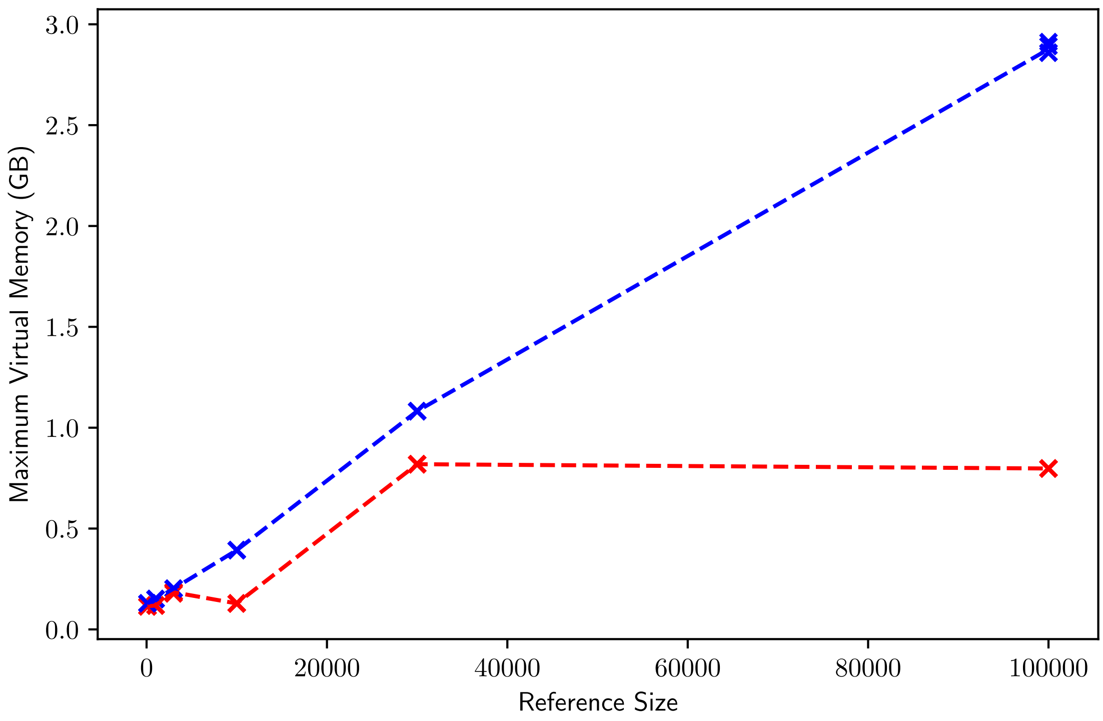


**Peak memory usage by GangSTR and ExpansionHunter vs. reference size** The x-axis shows the number of TRs in the reference set used. The y-axis shows maximum virtual memory usage in gigabytes. Lines give mean value across 5 runs. Points ("x") give raw data values for each of 5 runs. Red=GangSTR, blue=ExpansionHunter.

### Supplementary Figure 9

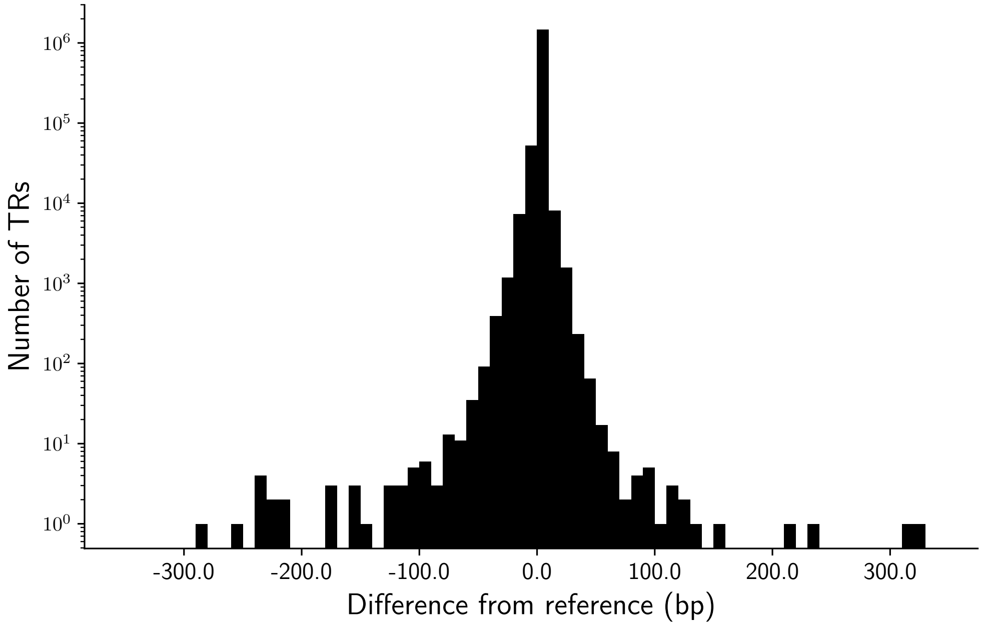


**Distribution of repeat lengths in NA12878 compared to the hg19 reference.** Y-axis is on a log10 scale.

### Supplementary Figure 10

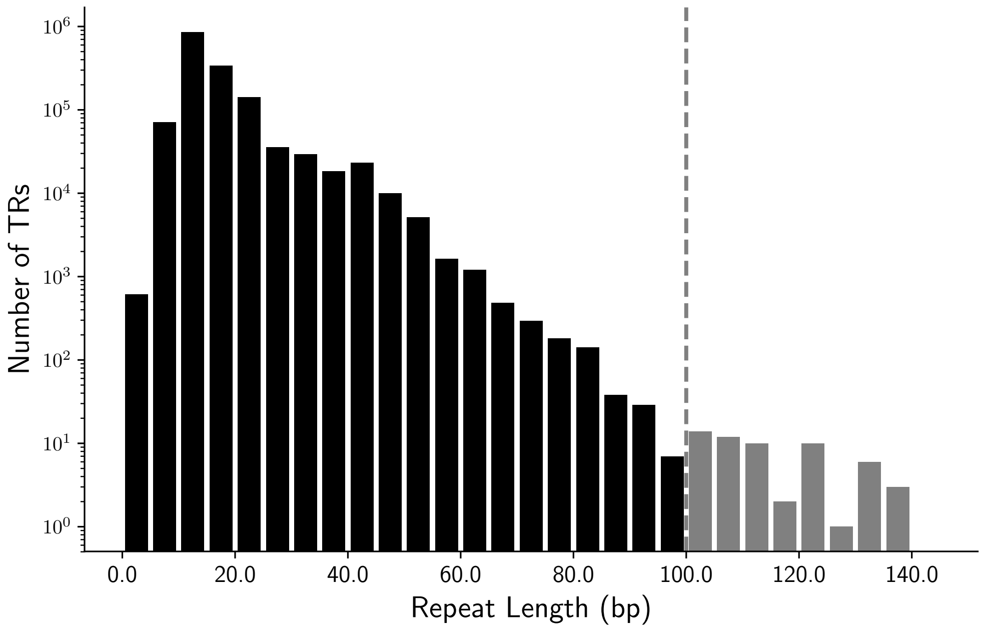


**Distribution of total repeat lengths in NA12878.** Y-axis is on a log10 scale. Gray bars to the right of the dashed line indicate alleles longer than the read length of 101bp.

### Supplementary Figure 11

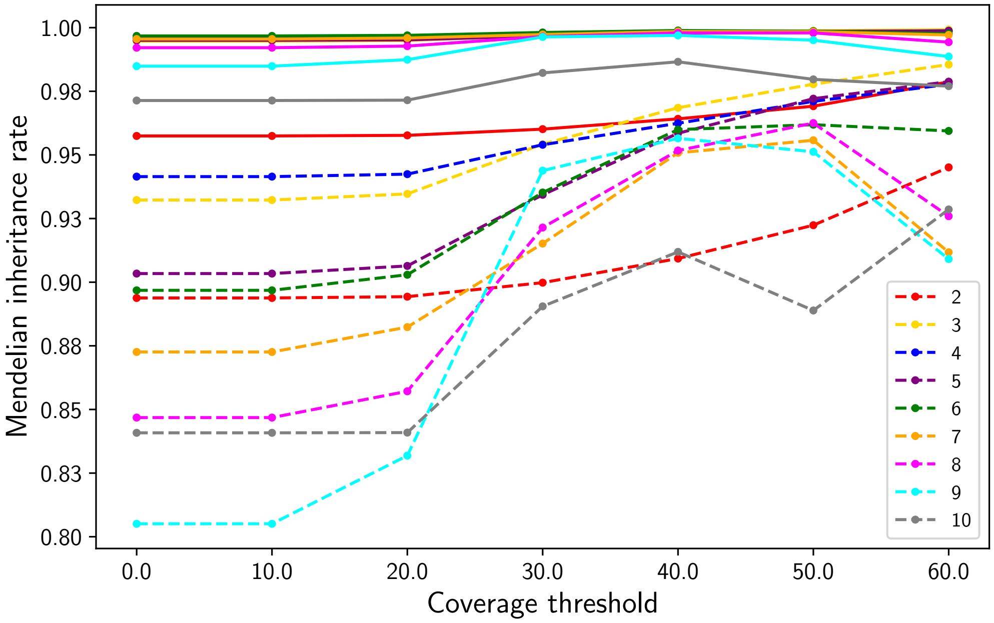


**Mendelian inheritance of GangSTR genotypes in a CEU trio as a function of the number of informative read pairs** Mendelian inheritance of GangSTR genotypes in a CEU trio as a function of the number of informative read pairs. Colors denote repeat lengths. Solid lines give mean Mendelian inheritance rate across all TRs, computed using maximum likelihood GangSTR genotypes as described in Methods. Dashed lines are computed after excluding loci where all three samples were homozygous for the reference allele.

### Supplementary Figure 12

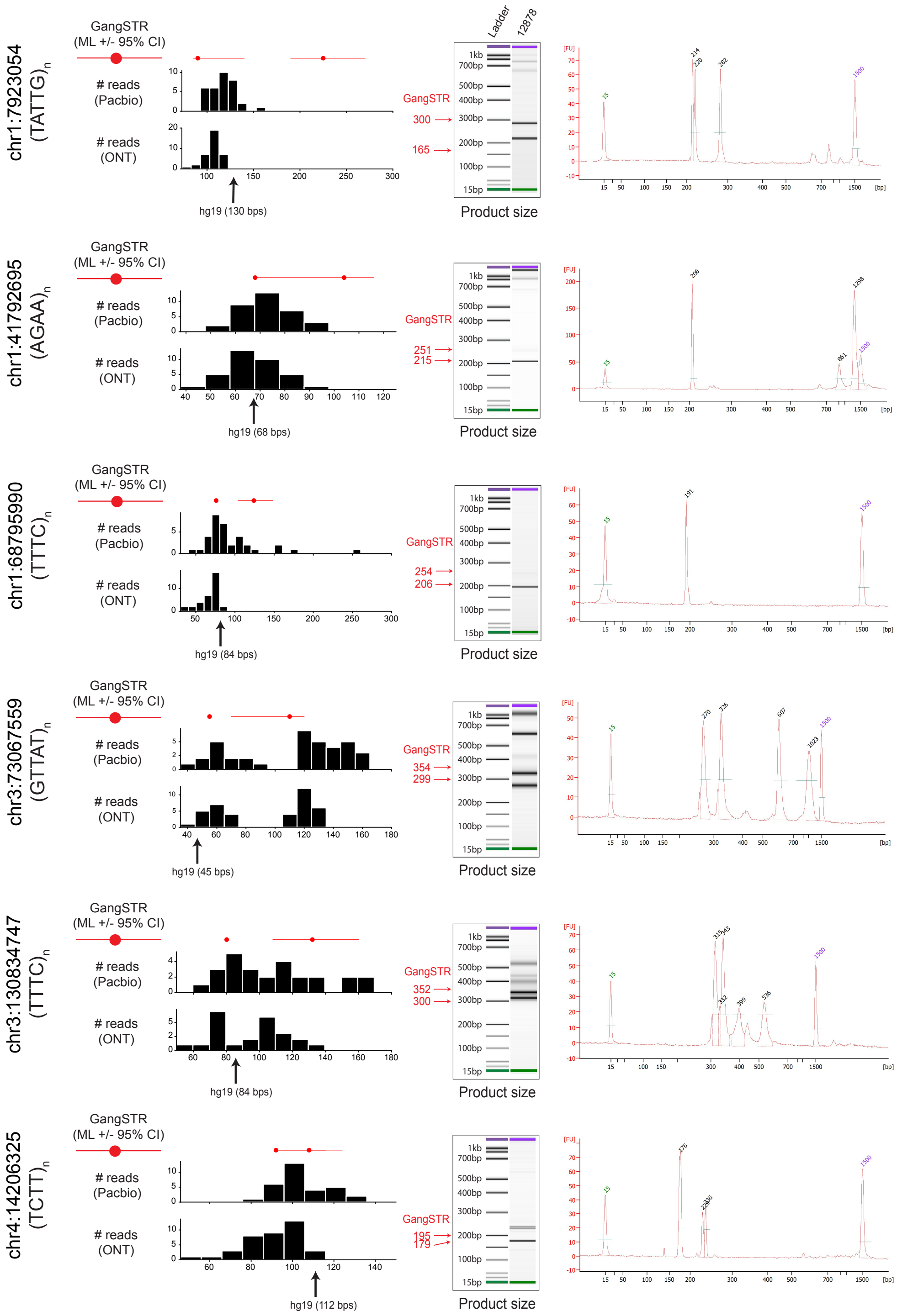


Continued on next page.

Continued from previous page.

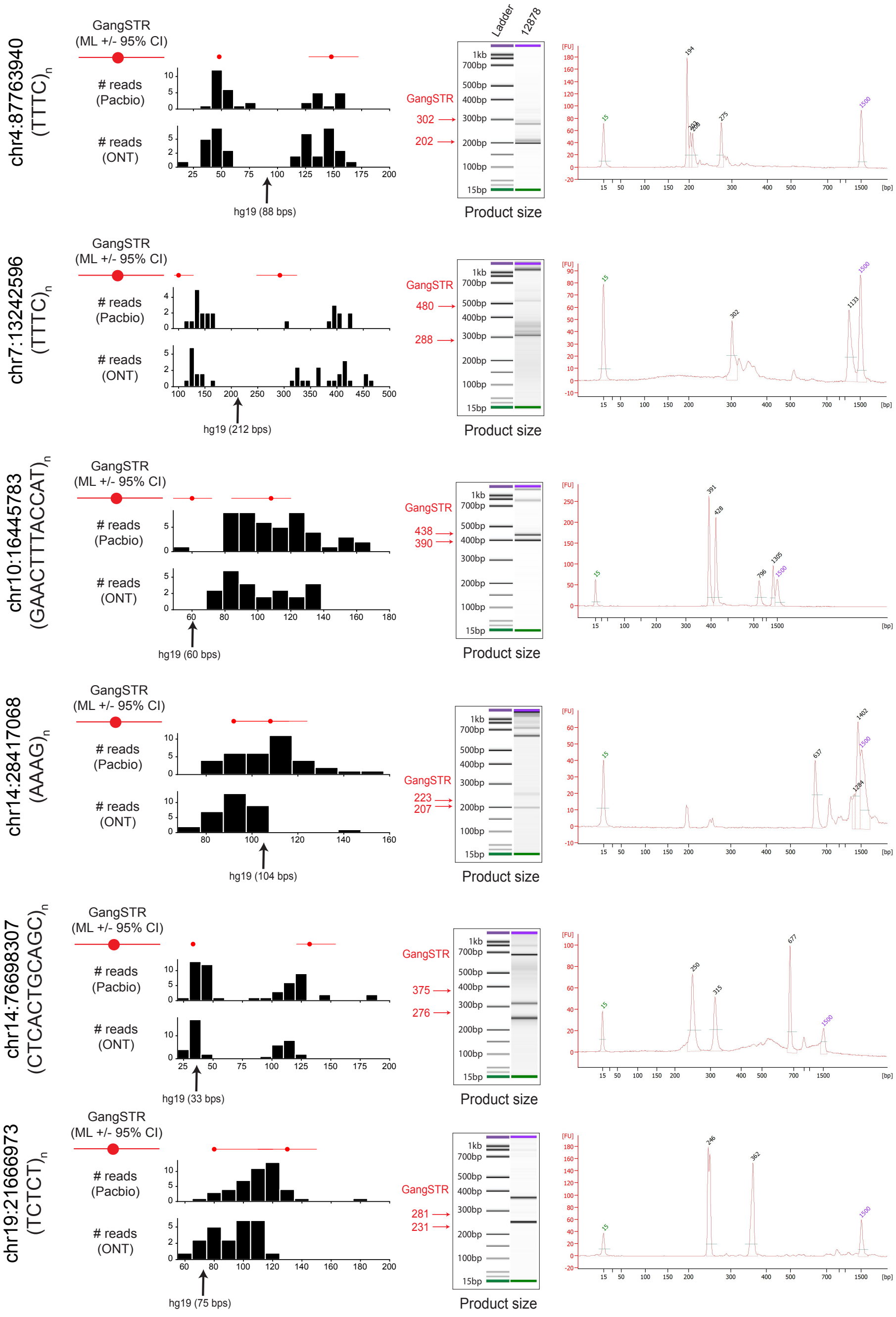

**Discovery and validation of genome-wide TR expansions.** For each of the 12 TRs shown, left plots compare GangSTR genotypes to those predicted by long reads. Red dots give the maximum likelihood repeat lengths predicted by GangSTR and red lines give the 95% confidence intervals for each allele. Black histograms give the distribution of repeat lengths supported by PacBio (top) and ONT (bottom) reads. The black arrow denotes the length in hg19. The middle plots show PCR product sizes for each TR as estimated using capillary electrophoresis. Left bands show the ladder and right bands show product sizes in NA12878. Green and purple bands show the lower and upper limits of the ladder, respectively. Red arrows and numbers give product sizes expected for the two alleles called by GangSTR. Right plots give the capillary electrophoresis traces produced by the Agilent Bioanalyzer.

### 3 Supplementary Tables

#### Supplementary Table 1: Target pathogenic repeats used in benchmarking experiments.

| Abbreviation | Disease | Gene | Motif | Repeat location | Pathogenic cutoff | Simulation repeat range[a] |
|---|---|---|---|---|---|---|
| SCA6 | Spinocerebellar ataxia 6 | *CACNA1A* | CAG | chr19:13207859-13207897 (hg38) chr19:13318673-13318711 (hg19) | 20 (60 bps) | [4, 7, 10, 13, 16, 19] |
| SCA2 | Spinocerebellar ataxia 2 | *ATXN2* | CAG | chr12:111598951-111599019 (hg38) chr12:112036755-112036823 (hg19) | 33 (99 bps) | [2, 8, 14, 20, 26, 32] |
| SCA7 | Spinocerebellar ataxia 7 | *ATXN7* | CAG | chr3:63912686-63912715 (hg38) chr3:63898362-63898391 (hg19) | 34 (102 bps) | [3, 9, 15, 21, 27, 33] |
| SCA1 | Spinocerebellar ataxia 1 | *ATXN1* | CAG | chr6:16327636-16327722 (hg38) chr6:16327867-16327953 (hg19) | 39 (117 bps) | [3, 10, 17, 24, 31, 38] |
| HTT | Huntingtons Disease | *HTT* | CAG | chr4:3074877-3074933 (hg38) chr4:3076604-3076660 (hg19) | 40 (120bps) | [4, 11, 18, 25, 32, 39] |
| SCA17 | Spinocerebellar ataxia 17 | *TBP* | CAG | chr6:170561908-170562021 (hg38) chr6:170870996-170871109 (hg19) | 43 (123 bps) | [2, 10, 18, 26, 34, 42] |
| DM1 | Myotonic Dystrophy 1 | *DMPK* | CTG | chr19:45770205-45770264 (hg38) chr19:46273463-46273522 (hg19) | 50 (150 bps) | [4, 13, 22, 31, 40, 49] |
| SCA12 | Spinocerebellar ataxia 12 | *PPP2R2B* | CAG | chr5:146878729-146878758 (hg38) chr5:146258292-146258321 (hg19) | 51 (153 bps) | [5, 14, 23, 32, 41, 50] |
| SCA3 | Spinocerebellar ataxia 3 | *ATXN3* | CAG | chr14:92071011-92071034 (hg38) chr14:92537355-92537378 (hg19) | 60 (120bps) | [4, 15, 26, 37, 48, 59] |
| C9ORF72 | ALS | *C9ORF72* | GGCCCC | chr9:27573529-27573546 (hg38) chr9:27573527-27573544 (hg19) | 31 (186 bps) | [5, 10, 15, 20, 25, 30] |
| SCA8 | Spinocerebellar ataxia 8 | *ATXN8OS* | CTG | chr13:70139384-70139428 (hg38) chr13:70713516-70713560 (hg19) | 80 (240 bps) | [4, 19, 34, 49, 64, 79] |
| FMR1 | Fragile X syndrome | *FMR1* | CGG | chrX:147912051-147912110 (hg38) chrX:146993569-146993628 (hg19) | 200 (600 bps) | [4, 43, 82, 121, 160, 199] |
| SCA36 | Spinocerebellar ataxia 36 | *NOP56* | GGCCTG | chr20:2652734-2652757 (hg38) chr20:2633380-2633403 (hg19) | 650 (3900 bps) | [4, 133, 262, 391, 520, 649] |
| SCA10 | Spinocerebellar ataxia 10 | *ATXN10* | ATTCT | chr22:45795355-45795424 (hg38) chr22:46191235-46191304 (hg19) | 800 (4000 bps) | [4, 163, 322, 481, 640, 799] |

[a] The other allele for all TRs covers the range (5, 1005) with step size 100. Simulation repeat range is given in terms of repeat copy number.

Repeat locations are given for both hg19 and hg38 genomic coordinates.

#### Supplementary Table 2: Candidate TRs long alleles (>101bp) in NA12878.

| Coord (hg19) | Refcopy[a] | Motif | GangSTR[b] | P(het)[c] | P(hom)[d] | MI[e] | Parent[f] | PacBio[g] | ONT[h] |
|---|---|---|---|---|---|---|---|---|---|
| chr1:2897558 | 4 | AACAGGAGGTCTGGT | 1,7 | 1.00 | 0.00 | True | NA | 147 | 108 |
| chr1:7923054 | 26 | AATAC | 18,45 | 0.93 | 0.07 | True | NA12891,NA12892 | 154 | 123 |
| chr1:22720748 | 13 | AAAG | 16,30 | 1.00 | 0.00 | True | NA12891,NA12892 | 163 | 87 |
| chr1:35267829 | 44 | AAACC | 20,90 | 0.28 | 0.72 | True | NA12891,NA12892 | 377 | 275 |
| chr1:41792695 | 17 | AAAG | 17,26 | 1.00 | 0.00 | True | NA12891 | 225 | 93 |
| chr1:61347419 | 3 | AAAG | 3,33 | 1.00 | 0.00 | True | NA12891,NA12892 | 64 | 23 |
| chr1:64329379 | 19 | AATAC | 17,22 | 1.00 | 0.00 | True | NA12892 | 231 | 109 |
| chr1:68795990 | 21 | AAAG | 19,31 | 1.00 | 0.00 | True | NA12892 | 363 | 85 |
| chr1:154098099 | 29 | AAGG | 26,26 | 0.74 | 0.26 | NA | NA12891 | 241 | 217 |
| chr1:208680308 | 6 | AATGTGGTATATATACAT | 4,5 | 0.67 | 0.29 | NA | NA | 168 | 141 |
| chr10:16445783 | 5 | AAAGTTCATGGT | 5,9 | 1.00 | 0.00 | True | NA12891,NA12892 | 169 | 135 |
| chr10:99438638 | 60 | AC | 46,59 | 0.73 | 0.27 | True | NA12891,NA12892 | 147 | 168 |
| chr10:125413213 | 15 | AAAGG | 14,21 | 1.00 | 0.00 | True | NA | 192 | 134 |
| chr11:17574076 | 3 | ACACAGGACAGGTGGGGG | 6,6 | 0.13 | 0.87 | NA | NA | 557 | 495 |
| chr11:31932832 | 26 | AAAGG | 20,49 | 0.58 | 0.42 | True | NA12891,NA12892 | 280 | 199 |
| chr11:107461059 | 53 | AG | 47,62 | 0.77 | 0.23 | True | NA12891,NA12892 | 223 | 89 |
| chr12:15314073 | 22 | AAAG | 22,28 | 0.98 | 0.02 | True | NA12891,NA12892 | 211 | 127 |
| chr12:117836405 | 20 | AAAAT | 9,21 | 1.00 | 0.00 | True | NA12891 | 138 | 111 |
| chr13:29027163 | 23 | AAAGG | 16,23 | 0.88 | 0.12 | True | NA12891,NA12892 | 205 | 105 |
| chr13:44716269 | 35 | AGCCG | 14,22 | 1.00 | 0.00 | True | NA12891 | 182 | 133 |
| chr13:87882390 | 3 | AAAG | 3,33 | 1.00 | 0.00 | True | NA12891,NA12892 | 92 | 60 |
| chr13:96047512 | 16 | AAAGG | 17,24 | 1.00 | 0.00 | True | NA12891 | 155 | 113 |
| chr14:28417068 | 26 | AAAG | 23,27 | 0.80 | 0.20 | True | NA12891,NA12892 | 216 | 143 |
| chr14:76698307 | 3 | ACTGCAGCCTC | 3,12 | 1.00 | 0.00 | True | NA12891,NA12892 | 535 | 125 |
| chr15:54367612 | 5 | AAGCTCCGGCTCACTGC | 4,5 | 0.88 | 0.11 | True | NA | 189 | 94 |
| chr15:61429219 | 20 | AAAG | 21,26 | 1.00 | 0.00 | True | NA12892 | 143 | 112 |
| chr15:90651456 | 13 | AAAAT | 8,21 | 1.00 | 0.00 | NA | NA | 114 | 119 |
| chr16:3899380 | 55 | AC | 36,55 | 0.97 | 0.03 | True | NA12891,NA12892 | 584 | 124 |
| chr16:10297534 | 29 | AAAG | 13,34 | 1.00 | 0.00 | True | NA12892 | 288 | 106 |
| chr16:23770703 | 10 | AAAG | 16,30 | 1.00 | 0.00 | True | NA12892 | 249 | 91 |
| chr16:50509578 | 53 | AAAG | 17,39 | 1.00 | 0.00 | True | NA12891,NA12892 | 169 | 90 |
| chr16:58865692 | 10 | AAGGAGGG | 7,14 | 1.00 | 0.00 | True | NA12891,NA12892 | 140 | 110 |
| chr17:32835083 | 21 | AAAGG | 18,27 | 0.92 | 0.08 | True | NA12891,NA12892 | 146 | 146 |
| chr19:21666973 | 15 | AAGAG | 16,26 | 0.95 | 0.05 | True | NA12891,NA12892 | 181 | 116 |
| chr19:39720793 | 15 | AAAGG | 20,26 | 0.55 | 0.45 | True | NA12891,NA12892 | 218 | 275 |
| chr2:54425083 | 18 | AATAC | 17,24 | 1.00 | 0.00 | True | NA12892 | 156 | 140 |
| chr2:163609414 | 25 | AAAAG | 67,89 | 0.00 | 1.00 | True | NA12891,NA12892 | 426 | 287 |
| chr2:178524872 | 3 | AAAG | 2,26 | 1.00 | 0.00 | True | NA12892 | 81 | 14 |
| chr21:36720944 | 18 | AATAG | 19,37 | 0.78 | 0.22 | True | NA12891,NA12892 | 182 | 156 |
| chr22:22928272 | 4 | AAAG | 4,32 | 1.00 | 0.00 | NA | NA12891 | 70 | 16 |
| chr22:22928365 | 28 | AAAG | 23,31 | 0.75 | 0.25 | NA | NA12891 | NA | NA |
| chr22:47769363 | 3 | AAGGGAGGCCAGGAGGAG | 3,6 | 1.00 | 0.00 | True | NA12891 | 117 | 113 |
| chr3:5830605 | 11 | AAATGCACAGGAAT | 6,16 | 0.61 | 0.39 | True | NA12891,NA12892 | 230 | 180 |
| chr3:43219613 | 18 | AAAG | 18,26 | 1.00 | 0.00 | True | NA12892 | 157 | 109 |
| chr3:73067559 | 9 | AACAT | 11,22 | 1.00 | 0.00 | True | NA12892 | 165 | 135 |
| chr3:86384908 | 19 | AAAAT | 18,22 | 0.70 | 0.30 | NA | NA | 162 | 120 |
| chr3:130834747 | 21 | AAAG | 20,33 | 1.00 | 0.00 | True | NA12892 | 167 | 135 |
| chr4:14206325 | 28 | AAAG | 23,27 | 0.90 | 0.10 | True | NA12891,NA12892 | 203 | 115 |
| chr4:21716410 | 61 | AAAAT | 18,48 | 0.71 | 0.29 | True | NA12891,NA12892 | 293 | 287 |
| chr4:58380837 | 3 | AAAG | 3,42 | 1.00 | 0.00 | True | NA12891,NA12892 | 50 | 23 |
| chr4:87763940 | 22 | AAAG | 12,37 | 1.00 | 0.00 | True | NA12892 | 303 | 168 |
| chr4:90302001 | 3 | AAAG | 3,26 | 1.00 | 0.00 | True | NA | 129 | 48 |
| chr5:15005273 | 3 | AAAG | 3,31 | 1.00 | 0.00 | True | NA12891,NA12892 | 45 | 15 |
| chr5:28661573 | 4 | AAAG | 4,26 | 1.00 | 0.00 | True | NA12891 | 68 | 27 |
| chr5:75792512 | 3 | AAAG | 3,28 | 1.00 | 0.00 | True | NA12891,NA12892 | 67 | 18 |
| chr5:157994659 | 20 | AAAG | 17,26 | 1.00 | 0.00 | True | NA12891 | 143 | 107 |
| chr6:128925487 | 18 | AGAGCGGG | 5,17 | 1.00 | 0.00 | True | NA12891,NA12892 | 515 | 161 |
| chr7:2852271 | 12 | ACATC | 18,39 | 0.78 | 0.22 | True | NA12891,NA12892 | 188 | 181 |
| chr7:6460939 | 18 | AGCGCGGGAGGCGCAGGC | 4,6 | 0.99 | 0.00 | True | NA12892 | 652 | 431 |
| chr7:13242596 | 53 | AAAG | 25,73 | 0.60 | 0.40 | True | NA12891,NA12892 | 428 | 469 |
| chr7:68065433 | 15 | AAAG | 16,31 | 1.00 | 0.00 | False | NA | 140 | 95 |
| chr7:80126869 | 8 | ACATATACGTATAT | 3,8 | 1.00 | 0.00 | True | NA12891,NA12892 | 495 | 165 |
| chr7:105084942 | 6 | AACACCTATAGC | 3,8 | 0.88 | 0.00 | True | NA | 544 | 318 |
| chr7:127898719 | 17 | AAAG | 22,26 | 0.89 | 0.11 | NA | NA12892 | 184 | 126 |
| chr7:134201476 | 15 | AAAAG | 15,22 | 1.00 | 0.00 | True | NA12891 | 156 | 115 |
| chr8:59629304 | 4 | ACATACATATATATGAT | 4,5 | 0.96 | 0.00 | NA | NA12892 | 226 | 114 |
| chr8:119927182 | 13 | AGAGAGCG | 11,20 | 0.90 | 0.10 | True | NA12891,NA12892 | 263 | 136 |
| chr8:130361920 | 15 | AAAAT | 16,21 | 0.88 | 0.12 | NA | NA12892 | 141 | 116 |
| chr8:140126207 | 5 | AAGACGACTCCACCCCACAG | 3,6 | 1.00 | 0.00 | NA | NA | 133 | 121 |

[a]Number of copies of the motif in hg19

[b]Maximum likelihood diploid repeat copy number returned by GangSTR.

[c]Posterior probability that the genotype is heterozygous for one allele greater than 101bp.

[d]Posterior probability that the genotype is homozygous for both alleles greater than 101bp.

[e]Indicates whether confidence intervals in the trio are consistent with Mendelian inheritance. NA indicates one or more parents failed filtering steps so Mendelian inheritance couldn't be determined.

[f]Lists which parents show evidence (>80% posterior probability) of an expansion

[g]Maximum allele length supported by PacBio reads for NA12878. "-" indicates no PacBio reads were found in the region.

[h]Maximum allele length supported by ONT reads for NA12878. "-" indicates no ONT reads were found in the region.

**Supplementary Table 3: Enrichment of motifs with long alleles in NA12878**.

| Motif | Num. TRs | P-val |
|---|---|---|
| AAAG | 27 | 2.675828e-19 |
| AAAGG | 6 | 2.179721e-07 |
| AATAC | 3 | 2.835557e-05 |
| AAAAT | 5 | 1.739327e-01 |
| AAAAG | 2 | 5.173664e-01 |
| AC | 2 | 9.759742e-01 |

All motifs found at least twice in long alleles (>101bp) in NA12878 are shown. P-values were computed using a one-sided Fisher's exact test.

#### Supplementary Table 4: Comparison of STRetch and GangSTR output.

| Chrom | STR Pos (hg19) | Motif | hg19 copy num | STRetch copy num [a] | GangSTR gt [b] | GangSTR filter [c] | QEXP [d] |
|---|---|---|---|---|---|---|---|
| chr1 | 41410562 | AAAGC | 10 | 11.80 | . | SpanBoundOnly | . |
| chr1 | 59446780 | AAAG | 41 | 43.30 | 15,17 | PASS | 1.00,0.00,0.00 |
| chr1 | 85576133 | AAAT | 7 | 9.30 | 11,13 | PASS | 1.00,0.00,0.00 |
| chr1 | 208440810 | AGC | 8 | 14.30 | 9,22 | PASS | 1.00,0.00,0.00 |
| chr1 | 234275697 | AAAAT | 9 | 10.80 | 14,17 | PASS | 0.89,0.11,0.00 |
| chr10 | 49500011 | AAAG | 5 | 8.50 | 7,22 | PASS | 0.87,0.13,0.00 |
| chr10 | 99438637 | AC | 59 | 63.60 | 46,59 | PASS | 0.00,0.73,0.27 |
| chr10 | 106199462 | AAC | 8 | 11.10 | 14,15 | PASS | 1.00,0.00,0.00 |
| chr11 | 62855843 | AACATC | 3 | 4.50 | 8,8 | PASS | 1.00,0.00,0.00 |
| chr12 | 79152282 | AATGT | 11 | 12.80 | 11,12 | PASS | 1.00,0.00,0.00 |
| chr13 | 113588869 | AGC | 7 | 10.10 | 9,17 | PASS | 1.00,0.00,0.00 |
| chr15 | 75186380 | AC | 33 | 37.60 | . | LowCallDepth, SpanBoundOnly | . |
| chr16 | 17564764 | CCG | 4 | 8.70 | . | LowCallDepth, SpanBoundOnly | . |
| chr16 | 57893926 | AAAAT | 13 | 16.80 | 15,15 | PASS | 1.00,0.00,0.00 |
| chr16 | 65292358 | ACATAT | 3 | 6.10 | 4,12 | PASS | 1.00,0.00,0.00 |
| chr17 | 15378610 | AAAAG | 9 | 11.80 | 13,16 | PASS | 1.00,0.00,0.00 |
| chr18 | 4057004 | AC | 27 | 31.60 | 24,28 | PASS | 1.00,0.00,0.00 |
| chr19 | 53608032 | AACAT | 4 | 5.80 | 5,10 | PASS | 1.00,0.00,0.00 |
| chr2 | 17512498 | ACACAT | 3 | 5.30 | 5,10 | PASS | 1.00,0.00,0.00 |
| chr2 | 112925446 | AACAT | 9 | 12.80 | . | SpanBoundOnly | . |
| chr2 | 163609413 | AAAAG | 24 | 25.80 | 67,89 | PASS | 0.00,0.00,1.00 |
| chr2 | 206247807 | AGAT | 13 | 15.30 | 14,22 | PASS | 0.89,0.11,0.00 |
| chr20 | 49282720 | AAAAG | 7 | 23.90 | . | NOCALL | . |
| chr21 | 34315567 | AAAAG | 7 | 8.80 | 9,14 | PASS | 1.00,0.00,0.00 |
| chr22 | 22928364 | AAAG | 27 | 29.30 | 23,31 | PASS | 0.00,0.75,0.25 |
| chr3 | 47788930 | AAATAT | 6 | 12.40 | 10,13 | PASS | 1.00,0.00,0.00 |
| chr3 | 121505017 | AAAAG | 8 | 24.90 | 13,13 | PASS | 1.00,0.00,0.00 |
| chr3 | 123773204 | AAAAAT | 3 | 7.80 | 8,9 | PASS | 1.00,0.00,0.00 |
| chr3 | 139970426 | AAAGG | 10 | 11.80 | 13,17 | PASS | 0.92,0.08,0.00 |
| chr3 | 188449305 | AAAAC | 3 | 5.80 | 4,12 | PASS | 1.00,0.00,0.00 |
| chr4 | 10768264 | AAAAG | 14 | 17.80 | . | SpanBoundOnly | . |
| chr4 | 30718515 | AGC | 11 | 20.60 | 12,28 | PASS | 1.00,0.00,0.00 |
| chr4 | 36460354 | ACACAT | 7 | 17.70 | 12,12 | PASS | 0.98,0.02,0.00 |
| chr5 | 11846 | ACCCCG | 3 | 5.30 | . | LowCallDepth | . |
| chr5 | 168182394 | AAAG | 13 | 15.30 | 13,13 | PASS | 1.00,0.00,0.00 |
| chr5 | 176523778 | AGG | 3 | 7.70 | 4,4 | PASS | 1.00,0.00,0.00 |
| chr6 | 43120472 | AACAT | 11 | 12.80 | 11,16 | PASS | 1.00,0.00,0.00 |
| chr6 | 65241442 | AAAAT | 9 | 11.80 | 9,15 | PASS | 1.00,0.00,0.00 |
| chr6 | 72287642 | AGAGAT | 3 | 12.80 | . | SpanBoundOnly | . |
| chr7 | 2852270 | ACATC | 11 | 23.80 | 18,39 | PASS | 0.00,0.78,0.22 |
| chr7 | 55955293 | CCG | 12 | 15.10 | 9,20 | PASS | 1.00,0.00,0.00 |
| chr7 | 105372194 | AAAAG | 10 | 11.80 | 11,17 | PASS | 0.94,0.06,0.00 |
| chr7 | 134201475 | AAAAG | 14 | 16.80 | 15,22 | PASS | 0.00,1.00,0.00 |
| chr8 | 87100322 | AAAAT | 6 | 8.80 | 9,14 | PASS | 1.00,0.00,0.00 |
| chr8 | 136094609 | ATCCC | 5 | 6.80 | 6,12 | PASS | 1.00,0.00,0.00 |

[a] Estimated repeat copy number returned by STRetch

[b] Maximum likelihood genotype (in terms of repeat copy number) returned by GangSTR.

[c] Locus-level GangSTR filters

[d] Expansion probability returned by GangSTR, which gives the probability of no expansion, a heterozygous expansion, or a homozygous expansion based on comparison to a predefined threshold. In this case the threshold was set to the read length of 101bp.

#### Supplementary Table 5: Primers for capillary electrophoresis validation.

| Chrom | STR Pos (hg19) | Forward Primer | Reverse Primer |
|---|---|---|---|
| chr1 | 7923054 | CAATAAGGCCTACCCTGACG | GGGCAACAAGAGCAAAACTT |
| chr1 | 41792695 | AGCTGCTTGAGAAGCTGAGG | CCCCATGGCTTTAACTCACT |
| chr1 | 68795990 | TTCCTCTCCCCAACACTTTTT | TGAGCCTCAGGAGATTGTTG |
| chr3 | 73067559 | GGTTGACAGCGGGATTTAAG | GAGCCATGGACACATCACTG |
| chr3 | 130834747 | TGGCGAGGTATTGTGGTAGA | TGACGAGTTAATGGGTGCAG |
| chr4 | 14206325 | ACAAACTTCTATGGGCTCGAT | CCTGGGCAAAGAGAGTGAAA |
| chr4 | 87763940 | AGCTGTCCTGAGTTGCATCA | GACTGAGGCAGGAGAAATGC |
| chr7 | 13242596 | GCATTTTCCTGATGGCTAAA | TTAGCCGGGTGTGGTAGC |
| chr10 | 16445783 | TGCCCAATAAGTATGAGAAGAACA | AAGTTCAAAAGGCCAGACCA |
| chr14 | 28417068 | CTGGGCGATAGAGCAAGACT | CCCTCATACCAAAGTGAACAAA |
| chr14 | 76698307 | ATAGAGTGCAGTGGGGCAAA | GAGCCCAAGAGTTCAACACC |
| chr19 | 21666973 | AGTACCGCTTAGGTCCAGCA | GGCAACAAGAGCGAAAACTC |
